# Induction of senescence renders cancer cells highly immunogenic

**DOI:** 10.1101/2022.06.05.494912

**Authors:** Inés Marín, Olga Boix, Andrea García, Isabelle Sirois, Adrià Caballe, Eduardo Zarzuela, Irene Ruano, Camille Stephan-Otto Attolini, Neus Prats, José Alberto López-Domínguez, Marta Kovatcheva, Elena Garralda, Javier Muñoz, Etienne Caron, María Abad, Alena Gros, Federico Pietrocola, Manuel Serrano

**Author notes:** Equal contribution*: O.B., A.G, I.S.

## Abstract

Cellular senescence is a stress response that activates innate immunity. However, the interplay between senescent cells and the adaptive immune system remains largely unexplored. Here, we show that senescent cells display enhanced MHC class I (MHC-I) antigen processing and presentation. Immunization of mice with senescent syngeneic fibroblasts generates CD8 T cells reactive against both normal and senescent fibroblasts, some of them targeting senescence-associated MHC-I-peptides. In the context of cancer, we demonstrate that senescent cancer cells trigger strong anti-tumor protection mediated by antigen-presenting cells and CD8 T cells. This response is superior to the protection elicited by cells undergoing immunogenic cell death. Finally, induction of senescence in patient-derived cancer cells exacerbates the activation of autologous tumor-reactive CD8 tumor-infiltrating lymphocytes (TILs) with no effect on non-reactive TILs. Our study indicates that immunization with senescent cancer cells strongly activates anti-tumor immunity, and this can be exploited for cancer therapy.

**STATEMENT OF SIGNIFICANCE:** Our study shows that senescent cells are endowed with a high immunogenic potential, superior to the gold standard of immunogenic cell death. The induction of senescence in cancer cells can be exploited to develop efficient and protective CD8-dependent anti-tumor immune responses.

## INTRODUCTION

Cellular senescence is a stress response that aims to eliminate unwanted cells (1). This response consists of a stable proliferative arrest together with the development of a vigorous proinflammatory secretome (2,3). Senescent cells recruit immune cells through their secretome to promote their own immune surveillance, thereby restoring tissue homeostasis (4). Immune clearance of senescent cells occurs in a context-dependent manner and is mediated by different populations of leukocytes, most of which belong to the innate immune system (4–6). Macrophages (7–10) and natural killer (NK) cells (10–16) have been described as the main cell types responsible for the elimination of senescent cells. Other components of the innate immune system, such as NKT cells (17), neutrophils (18), and natural IgM antibodies (19) have also been shown to participate in the elimination of senescent cells.

Regarding the adaptive immune system, it has been reported that senescent hepatocytes expressing oncogenic mutant *Nras* activate CD4 T cells, but not CD8 T cells, and this triggers macrophage-mediated elimination of the senescent hepatocytes (7). In this experimental system, CD4 T activation was found to be mediated by antigen-presenting cells and senescent hepatocytes expressing MHC-II (7). MHC-II expression has also been observed in senescent melanocytes but not in senescent keratinocytes or fibroblasts (20). The role of CD8 T cells in the elimination of senescent cells has been described through an antigen-independent mechanism. Specifically, senescent human skin fibroblasts upregulate HLA-E, which is an inhibitory signal for NK and CD8 T cells, and depletion of HLA-E renders senescent cells susceptible to elimination by both NK and CD8 T cells (14). In this study, we explored the immunogenic properties of senescent cells and their capacity to activate antigen-dependent CD8 T cell responses. We exploited these properties in the context of cancer, showing that senescent cancer cells are very efficient in triggering anti-tumor immune responses.

## RESULTS

### Senescent cells upregulate MHC class I antigen presentation

To identify proteins with potential immune regulatory activity specific to senescent cells, we performed a proteomic screen for plasma membrane-enriched fractions from senescent cells (SenC). We analyzed four cell types: two diploid primary fibroblasts (human IMR-90 fibroblasts and mouse embryonic fibroblasts [MEFs]) and two cancer cell lines (human melanoma SKMEL-103 and mouse melanoma B16-F10 [B16F10]) exposed to various senescence-inducing stimuli, namely, doxorubicin (doxo), the CDK4/6-inhibitor palbociclib (palbo), and the p53-activator nutlin-3A (nutlin), totaling seven senescence conditions (**Fig. 1A**). Successful induction of senescence was confirmed by monitoring senescence-associated beta-galactosidase (SABG) activity and the mRNA expression of the senescence-associated gene *CDKN1A*/*Cdkn1a* (**Supplementary Fig. S1A and S1B**). The “surfaceome” of the 7 senescent and the 4 non-senescent samples was obtained by mass spectrometry, each sample in biological triplicate (**Supplementary Table S1**). We used gene ontology (GO) to analyze proteins that were significantly increased in four or more senescence conditions as compared to their corresponding non-senescent controls and found that “*Antigen processing and presentation of antigen via MHC class I*” was the top enriched category (**Fig. 1B**) (**Supplementary Table S2**). Increased expression of classical MHC class I (MHC-I) molecules in senescent cells was confirmed by flow cytometry of mouse H-2K^b^/D^b^ in MEFs and B16F10 cells (**Fig. 1C and D**) as well as HLA-A/B/C in IMR-90 and SKMEL-103 cell lines (**Supplementary Fig. S1C**). In addition, we observed augmented levels of the non-classical MHC-I molecules H2-Qa1 and H2-Qa2 in senescent MEFs (**Supplementary Fig. S1D**).

**Figure 1.**
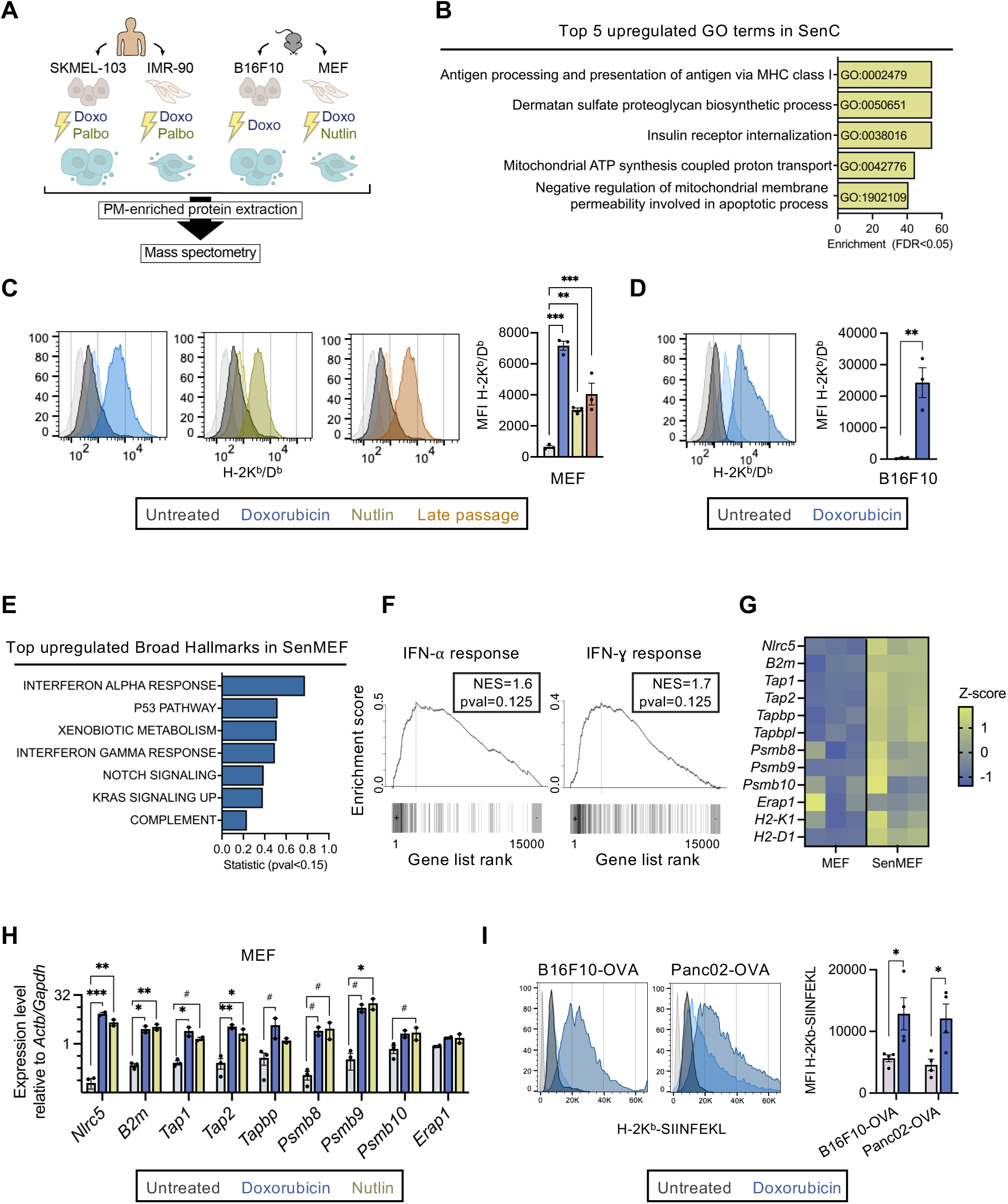
Senescent cells upregulate MHC class I antigen presentation. A. Schematics of the proteomic screen of the plasma membrane (PM)-enriched fraction of human (SKMEL-103, IMR-90) and murine (B16F10, mouse embryonic fibroblast [MEF]) cells, untreated or exposed to various senescence-inducing stimuli (doxo - doxorubicin, palbo - palbociblib and nutlin - nutlin-3a). 3 independent biological replicates per cell line were analyzed. B. Top 5 GO terms enriched in the proteins found upregulated in the plasma membrane fraction of senescent cells (SenC) (in 4 or more conditions of senescence, with a fold-change>1.5, FDR<0.05). C. Flow cytometry analysis of H-2K^b^/D^b^ expression in untreated *versus* senescent MEFs, in which senescence was induced by doxorubicin, nutlin or by late passaging. Representative histograms (left panel) showing the fluorescence signal of each stained sample and its unstained control (lighter color) and quantification after autofluorescence subtraction (right panel) of *n*=3 independent experiments are shown. ***p<0.001; **p<0.01, one-way ANOVA, compared to untreated MEFs. D. Flow cytometry analysis of H-2K^b^/D^b^ expression in untreated or senescent B16F10 cells, treated with doxorubicin. Representative histograms (left panel) showing the fluorescence signal of each stained sample and its unstained control (lighter color) and quantification after autofluorescence subtraction (right panel) of *n*=3 independent experiments are shown. **p<0.01, unpaired Student’s t test, compared to untreated cells. E. Top upregulated Broad Hallmarks from the differential expression analysis (RNAseq) of senescent MEFs, in which senescence was induced by doxorubicin compared to untreated cells. *n=*3 independent biological replicates were analyzed. F. Gene set enrichment analysis (GSEA) of IFN-⍺ and IFN-ɣ response (Broad Hallmarks) genes found upregulated in senescent MEFs compared to untreated cells. G. Normalized expression levels of antigen presentation machinery- and immunoproteasome-related genes from the RNA-seq analysis of untreated *versus* senescent MEFs. H. mRNA expression levels of antigen presentation machinery- and immunoproteasome-related genes in untreated *versus* senescent MEFs measured by qRT-PCR (relative to the average expression of housekeeping genes *Actb* and *Gapdh*). *n*=2 independent experiments. *p<0.05, ^#^p<0.1; one-way ANOVA, compared to untreated MEFs. I. Flow cytometry analysis of ovalbumin (OVA)-derived SIINFEKL peptide bound to H-2Kb presentation in untreated or senescent B16F10 and Panc02 cell lines, stably expressing OVA. Representative histograms (left panel) showing the fluorescence signal of each stained sample and its unstained control (lighter color) and quantification after autofluorescence subtraction (right panel) are shown. *p<0.05 unpaired Student’s t test, compared to untreated cells.

The expression of MHC-I presentation machinery is upregulated by the type I and type II interferon (IFN) pathways (21), and elevated IFN signaling has been previously described as a molecular feature of cellular senescence (22). In line with these findings, transcriptomic analysis of doxo-senescent MEFs, doxo-senescent murine pancreatic cancer Panc02 cells, and palbo-senescent SKMEL-103 cells revealed upregulation of IFN transcriptional signatures compared to their non-senescent counterparts (**Fig. 1E and F and Supplementary Fig. S1E-S1J**). Accordingly, the main gene components necessary for antigen processing and presentation through MHC-I were also found to be overexpressed in senescent cells in the transcriptomic analysis (**Fig. 1G and Supplementary Fig. S1G and S1J**). qRT-PCR analysis of these genes confirmed their upregulation in senescent cells, including the master transcriptional regulator of MHC-I-dependent immune responses, *Nlrc5* (23), and the main components of the immunoproteasome, such as *Psmb8*, *Psmb9* and *Psmb10* (**Fig. 1H and Supplementary Fig. S1K and S1L**). To test whether the transcriptional upregulation of the MHC-I-associated machinery was mirrored by an elevated presentation of antigens, we analyzed the expression of the H-2K^b^ restricted ovalbumin (OVA)-derived peptide SIINFEKL in senescent B16F10 and Panc02 cancer cells stably expressing OVA. We found that senescent cells presented more SIINFEKL bound to H-2K^b^ than their proliferating counterparts did (**Fig. 1I**). Taken together, these data support the concept that enhanced MHC-I antigen processing and presentation are a general feature of senescent cells.

### Senescent cells stimulate CD8 T cells

It is well established that stressed cells process and present non-mutated self-peptides that are not presented by non-stressed cells and can evoke an immune response (24–28). Considering that senescence is indeed a cellular response to stress, and in view of the enhanced capacity of senescent cells to process and present antigens, we wondered whether non-cancer senescent cells could induce an antigen-dependent immune response *in vivo*. To this end, we immunized mice with syngeneic primary fibroblasts, either untreated (MEF) or senescent (senMEFs). We also used vehicle with no cells as a negative control for immunization and ovalbumin (OVA) as a positive control for antigen-dependent activation against a known antigen (SIINFEKL) (**Fig. 2A**). All immunizations were performed using an immune adjuvant (CpG). After immunization, splenocytes from immunized animals were isolated, and the number of activated T cells was measured by IFN-γ enzyme-linked immunospot (ELISpot) in the absence of *ex vivo* stimuli (basal activation), or in the presence of MEF, senMEF, or SIINFEKL. Notably, basal activation was significantly higher in mice immunized with senMEF than under all other conditions **(Supplementary Fig. S2A**). Furthermore, splenocytes from senMEF-immunized mice responded strongly to *ex vivo* re-exposure to senMEF, yielding a number of spots over basal activity comparable to that of OVA-immunized splenocytes exposed to SIINFEKL (**Fig. 2B**). Mice immunized with senMEF also responded to MEF *ex vivo* although not as strongly as those immunized with senMEF (**Fig. 2B**). Moreover, CD8, but not CD4 T cells, derived from senMEF-immunized animals exhibited elevated levels of activation as monitored by flow cytometric detection of the activation markers CD69 and CD25 after *ex vivo* exposure to senMEF compared to normal MEF (**Fig. 2C, Supplementary Fig. S2B**). As a positive control, maximal possible activation of CD8 and CD4 T cells was achieved by treatment with a phorbol ester and calcium ionophore (abbreviated PMA+I) (**Supplementary Fig. S2C**). These observations suggest that immunization with senescent cells induces a CD8-dependent immune response.

**Figure 2:**
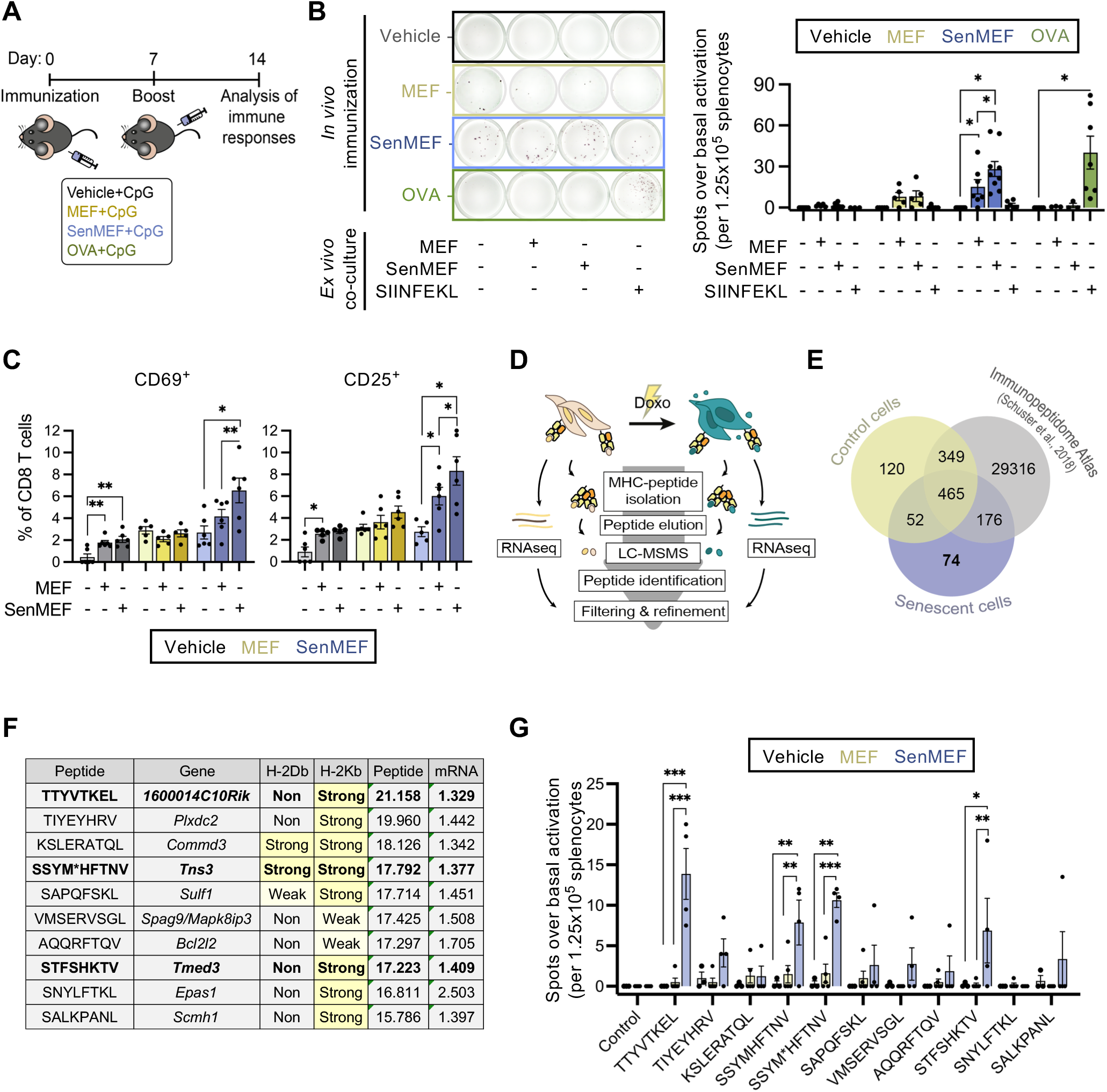
Senescent cells induce an adaptive immune response *in vivo* and present an altered immunopeptidome. A. Schematic outline of the immunization protocol used in this study. Briefly, immunocompetent C57BL/6J animals were subcutaneously immunized on days 0 and 7 with vehicle (no cell), untreated or senescent syngeneic fibroblasts (MEF or senMEF respectively) or ovalbumin (OVA), all done concomitantly with an immune adjuvant (CpG). One week later, animals were sacrificed and immune responses were tested *ex vivo*. B. ELISpot assay to detect IFN-γ production in splenocytes isolated from non-immunized mice or animals immunized with MEF, senMEF or OVA (*n=*3-7 mice per group). Splenocytes were cultured in RPMI, either alone (control) or co-cultured with untreated MEF (1:10 target-to-splenocyte ratio), senMEF (1:10 target-to-splenocyte ratio), or with SIINFEKL OVA-derived peptide. Restimulation of splenocytes isolated from OVA-immunized animals with the SIINKEFL peptide was used as a positive control. The number of spots for each condition above the control condition (background) was quantified. Representative picture (left panel) and quantification (right panel) are shown. *p<0.05; unpaired Student’s t test, compared to RPMI alone treatment. C. Measurement of CD8 T cell activation, as monitored via CD69 expression (left panel) or CD25 expression (right panel) in naïve *versus* MEF or senMEF-immunized animals, after co-culture with MEF or senMEF *ex vivo* (*n*=5-7 mice per group). D. Layout of combined immunopeptidomics and RNAseq analyses in untreated *versus* senescent MEFs. E. Venn diagram displaying peptides identified in control cells, senescent cells and the Mouse Immunopeptidome Atlas dataset (Schuster et al., 2018). F. List of selected peptides presented exclusively on senescent cells MHC-I together with their corresponding coding gene and their predicted binding to H-2K^b^ and H-2D^b^ (NetMHCpan v4.1). Levels of peptide obtained in the immunopeptidomic analysis and fold change expression the corresponding gene (senescent MEF *versus* untreated MEF) from RNAseq transcriptomic analysis are indicated. M* indicates M(+15.99), oxidized methionine. G. Selected peptides validated using ELISpot assay to detect IFN-γ production in splenocytes isolated from non-immunized mice or animals immunized with MEF or senMEF (*n*=3-5 mice per group). Splenocytes were cultured in RPMI, either alone as negative control (control) or supplemented with the different peptides selected from the immunopeptidome analysis as indicated. For SSYM*HFTNV peptide, both SSYMHFTNV and the modified SSYM*HFTNV were tested. The number of spots for each condition above the control condition (background) was quantified. ***p<0.001, **p<0.01, *p<0.05; unpaired Student’s t test, compared to vehicle or MEF immunization treatment.

### Senescence is associated to an altered immunopeptidome

To test whether CD8 T cell activation was directly linked to the presentation of immunogenic epitopes by senescent cells, we profiled the immunopeptidome of control and senescent MEF, in which senescence was induced by doxorubicin treatment (**Fig. 2D**). Mass spectrometry of peptides eluted from classical MHC-I identified 767 *bona fide* peptides corresponding to annotated proteins (see Methods). Interestingly, about 10% of these peptides (74), were detected in senescent cells but not in their non-senescent counterparts, nor in the “Mouse Immunopeptidome Atlas” (**Supplementary Table S3**). The latter consists of more than 30,000 MHC-I (H-2K^b^/D^b^) peptides obtained from 19 normal tissues and 4 murine cancer cell lines (29). As a criterion to prioritize the 74 senescence-associated MHC-I peptides, we selected those with upregulation (> 1.3-fold) of the coding mRNA in senescent *versus* untreated cells obtained by RNA-seq (10 out of 74) (**Fig. 2F**). To evaluate the immunogenic potential of these senescence-specific epitopes, we used individual peptides (of the 10 prioritized peptides) (**Fig. 2G**) or pooled combinations (of the remaining 64 peptides) (**Supplementary Fig. S2D and Table S3**) to re-stimulate splenocytes isolated from animals immunized with vehicle, MEF, or senMEF. As revealed by the IFN-γ ELISPOT assay, splenocytes isolated from senMEF-immunized animals were specifically activated by a subset of senescence-associated peptides (**Fig. 2G and Supplementary Fig. S2D**). In total, we identified 3 peptides from the set of 10 that were prioritized as immunogenic (**Fig. 2F**). Additionally, 3 peptide pools of the remaining set of peptides (up to 64) showed promising responses (**Supplementary Fig. 2D**). These results suggest that non-cancer senescent cells can elicit CD8 T cell responses against senescence-associated antigens, which further reinforces the notion that senescent cells are immunogenic.

### Senescent cancer cells display strong and sustained adjuvant properties

In addition to antigenic presentation, adaptive immune responses require the concomitant presence of adjuvant signals that promote the maturation of antigen-presenting cells (APCs), particularly dendritic cells (DCs) (30,31). Senescent cells secrete a plethora of proinflammatory cytokines and chemokines that recruit and activate immune cells (2,3), as well as damage-associated molecular patterns (DAMPs), such as calreticulin (CALR) (32). We compared the levels of DAMPs released between senescent B16F10 melanoma cells and cells undergoing immunogenic cell death (ICD). We chose ICD because it is the best-established method to boost tumor cell adjuvanticity, leading to efficient anti-tumor immunization (33,34). We observed that the levels of the prototypical DAMPs ATP and CALR in the conditioned media (CM) of ICD and senescent cells were similarly elevated compared to the untreated cells (**Fig. 3A and B and Supplementary Fig. S3A and S3B**).

**Figure 3:**
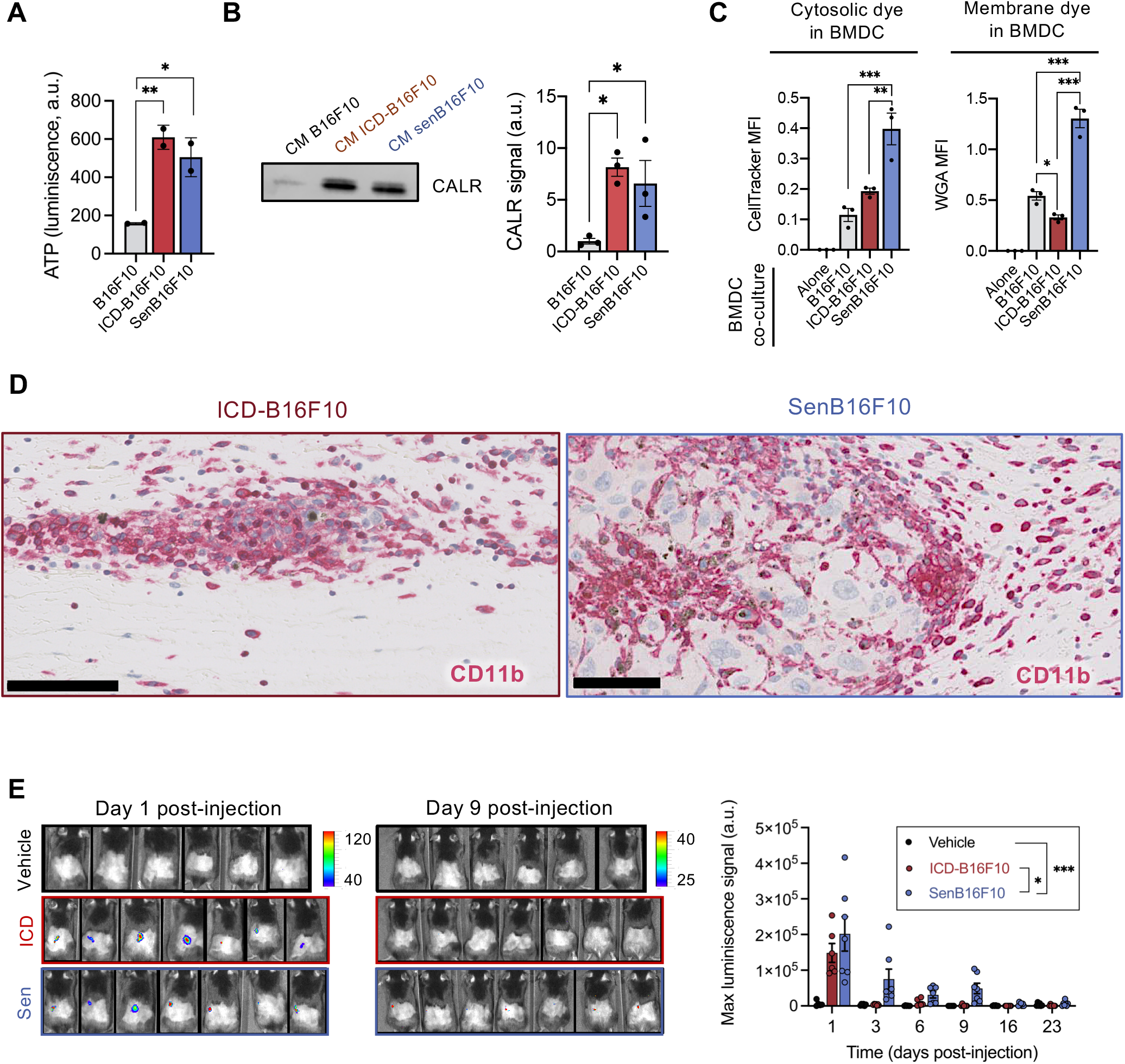
Senescent cancer cells display high immunogenic properties and sustained persistence *in vivo*. A. Levels of extracellular ATP in the conditioned media (CM) of 10^6^ untreated B16F10 cells, dying by immunogenic cell death induced by a high dose of doxorubicin (ICD-B16F10) and senescent B16F10 induced by a low dose of doxorubicin (senB16F10). *n*=2 independent experiments. **p<0.01 *p<0.05; one-way ANOVA test compared to untreated B16F10. B. Immunoblot detection of CALR in the CM of 10^6^ untreated B16F10, ICD-B16F10 and senB16F10. Representative image (left panel) and quantification (right panel) of *n*=3 independent experiments are shown. *p<0.05; one-way ANOVA test compared to untreated B16F10. C. Flow cytometry analysis of uptake of CFSE (cytosolic dye) or WGA-Alexa647 (membrane dye) by BMDCs from labeled untreated B16F10, ICD-B16F10 or senB10F10. Quantification after subtraction of autofluorescence from unstained BMDC of *n*=3 independent experiments. ***p< 0.001; **p<0.01 ; *p<0.05, one-way ANOVA test. D. Immunochemistry staining of CD11b^+^ cells (purple) in skin sections of animals 7 days after subcutaneous injection of ICD or senB16F10. Representative images selected by a histopathologist of *n*=5 animals per group are shown. Note that the brown pigmentation is due to the melanine. Scale bars for each images are shown (100µm). E. *In vivo* imaging detection of luciferase–expressing B16F10 treated with vehicle (*n*=6), ICD-B16F10 (*n*=7) or senB16F10 (*n*=7) at different time points after subcutaneous injection (as indicted). Representative images (left panels) and quantification (right panel) are shown. *p<0.05; Two-way ANOVA test compared to vehicle-treated group.

Upon recruitment and activation of immune cells, antigen capture is the next essential step in triggering an immune response (30,31). Therefore, we evaluated the efficiency of DCs capture of cytosolic and membrane antigens from senescent cells. Untreated, ICD, or senescent cancer cells were stained with membrane or cytosolic fluorescent dyes and co-cultured with syngeneic bone marrow-derived dendritic cells (BMDCs). We found that BMDCs (CD11c+) captured cytosolic and membrane dyes from B16F10 or Panc02 senescent cells more efficiently than from untreated or ICD cells (**Fig. 3C and Supplementary Fig. S3D**). As previously reported (35), ICD cells were efficient in delivering cytosolic dyes, but inefficient in delivering membrane dyes compared to untreated cells (**Fig. 3C and Supplementary Fig. S3D**).

To corroborate the adjuvant properties elicited by senescent cells *in vivo*, we injected mice subcutaneously with untreated, ICD, or senescent syngeneic B16F10 cancer cells and performed histological analysis. Interestingly, one week after injection, senescent melanoma cells were clearly visible in the subcutis as large pleomorphic and anaplastic cells, often containing granular melanin pigment, surrounded by abundant immune cells (CD45^+^), mainly of myeloid lineage (CD11b^+^) and some T cells (CD3^+^) (**Fig. 3D and Supplementary Fig. S3E and SEF**). Abundant immune and myeloid cells were also present at ICD injection sites, although, as expected, no evidence of viable tumor cells was found (**Fig. 3D and Supplementary Fig. S3E**). We wondered how long the senescent cells would persist at the injection site. To explore this, we injected luciferase-expressing senescent or ICD B16F10 cells and monitored their bioluminescence for 23 days. Because senescent cells are considerably larger than their non-senescent counterparts, we calculated the number of cells to be injected based on the total amount of protein. The luciferase signal from senescent cells was clearly detected 9 days post-injection but was undetectable on day 16 (**Fig. 3E**). In the case of ICD injection, the luciferase signal disappeared completely on day 3 (**Fig. 3E**). Collectively, these data support the notion that senescent cells provide prolonged and sustained immune recruitment, together with efficient delivery of cytosolic and membrane materials to dendritic cells.

### Immunization with senescent cancer cells promotes anti-cancer immune surveillance

Given that senescent cells exhibit enhanced antigenicity together with strong and persistent adjuvanticity, we postulated that senescent cancer cells could be used to promote an immune response against cancer. We first compared the prophylactic effect of immunization with senescent B16F10 melanoma cells (senB16F10) or cells undergoing ICD (ICD-B16F10) against subsequent tumor formation. To test the importance of cell viability in the case of senescent cells, we introduced another experimental condition that consisted of dying senescent cells by pre-treating senB16F10 cells with navitoclax, an agent known to induce apoptosis in senescent cells (*i.e.,* senolysis) (**Fig. 4A**). Immunizations were performed using equal amounts of protein for each condition. One week after immunization by subcutaneous injection, all experimental groups were re-challenged with 3×10^4^ viable B16F10 cells in the opposite flank. We observed that immunization with senB16F10 elicited superior anti-cancer protection compared to ICD-B16F10 or dying senB16F10. (**Fig. 4B and Supplementary Fig. S4A**). Similarly, senescent Panc02 (senPanc02) cells produced a stronger protective response than ICD-Panc02 cells (**Fig. 4C and Supplementary Fig. S4B**). The anti-tumor protection afforded by senescent cancer cells, B16F10 and Panc02, was reflected by a lower rate of tumor success and by a longer latency of those tumors that escaped immune control (in the case of Panc02, this was not measured because none of the animals developed tumors) (**Fig. 4B and C**s).

**Figure 4:**
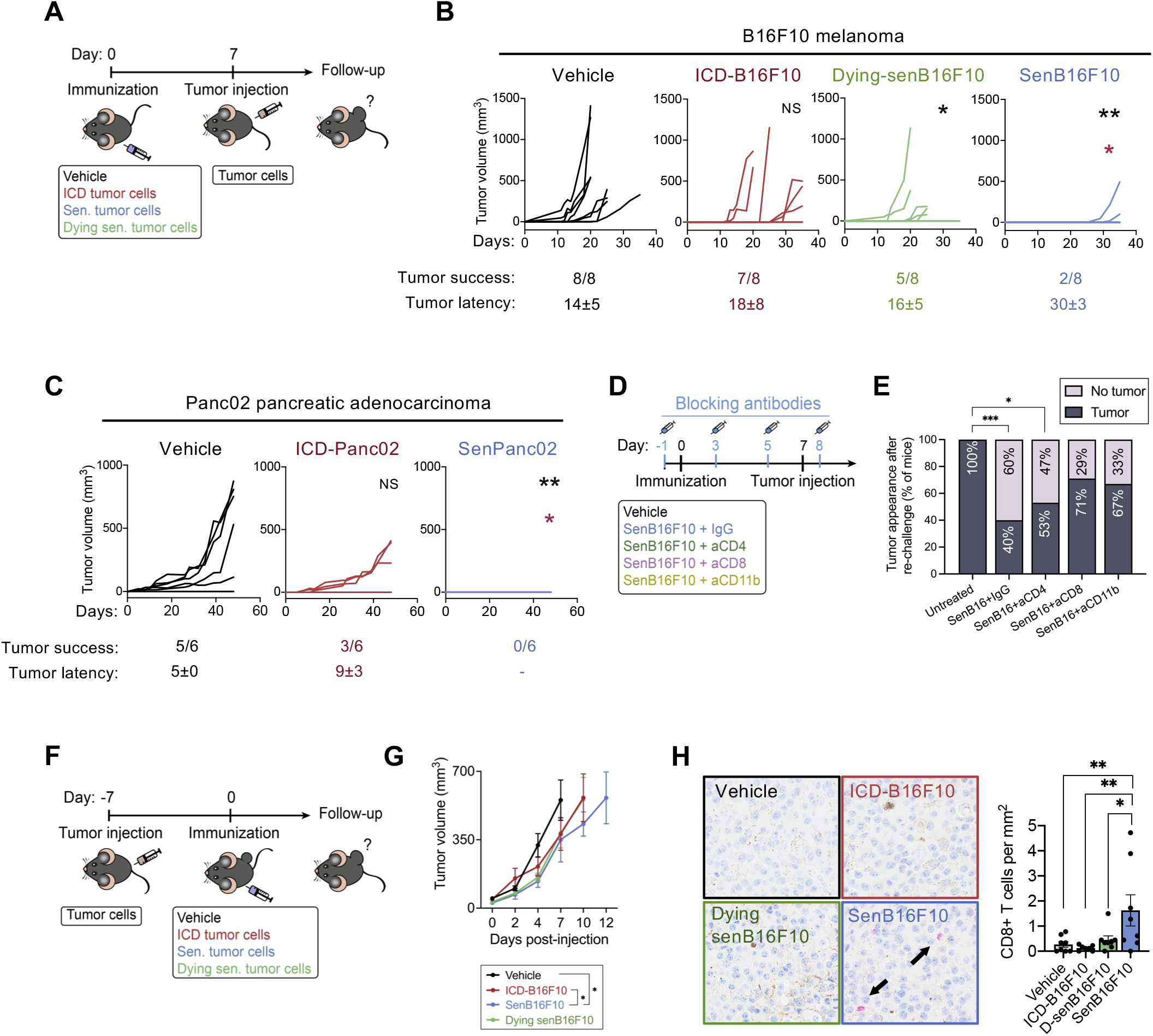
Immunization with senescent cancer cells promotes anti-cancer immune surveillance. A. Schematics of the cancer immunization protocol used in these studies. B. Individual tumor growth curves from vehicle-treated mice or mice immunized with B16F10 cells dying by immunogenic cell death induced by a high dose of doxorubicin (ICD-B16F10), senescent B16F10 induced by a low dose of doxorubicin (senB16F10) or senescent B16F10 cells dying by senolysis induced by navitoclax (dying-senB16F10) (*n*=8 mice per group). ***p<0.001, *p<0.05; Two-way ANOVA test compared to vehicle-treated (black) group. C. Individual tumor growth curves from vehicle-treated mice or mice immunized with Panc02 cells dying by immunogenic cell death induced by a high dose of doxorubicin (ICD-Panc02) or senescent Panc02 low dose of doxorubicin (senPanc02) (*n*=6 mice per group). ***p<0.001; Two-way ANOVA test compared to vehicle-treated (black) group. D. Scheme of the cancer immunization and immune depletion protocol used in this study. E. Tumor appearance after re-challenge in vehicle-treated mice (*n*=14) or mice immunized with senB16F10 treated with IgG (*n*=14) or the indicated blocking antibodies as described in d (*n*=15 for aCD4, *n*=14 for aCD8, or *n*=6 for aCD11b). ***p<0.001, *p<0.05, Fisher exact Test. F. Schematics of the therapeutic cancer immunization protocol used in these studies G. Grouped tumor growth of B16F10 tumor-bearing animals immunized with ICD-B16F10, dying senB16F10 or senB16F10. *p<0.05; Two-way ANOVA test (*n*=7-8) H. CD8 staining (purple) in B16F10 tumors from animals immunized with vehicle, ICD-B16F10 or senB16F10. Note that the brown pigmentation is due to the melanine. Representative pictures (left panels) and quantification (right panel) (*n*=7-8 mice per group). **p<0.01; one-way ANOVA test.

To gain a deeper understanding of the cancer preventive response induced by live senescent cancer cells, we depleted the major immune populations involved in adaptive anti-tumor immunity, namely CD4 and CD8 T cells and CD11b^+^ myeloid cells (**Fig. 4D and Supplementary Fig. S4C**). We found that depletion of CD8 T cells or CD11b^+^ myeloid cells significantly increased the rate of tumor formation, whereas depletion of CD4 T cells had a modest effect (**Fig. 4E and Supplementary Fig. S4D and S4E**).

We also tested whether immunization with senescent cells could inhibit tumor growth in mice already bearing tumors at the time of immunization (**Fig. 4F**). We found that animals immunized with senescent cells showed a moderate but significantly reduced rate of tumor growth compared with animals immunized with ICD cells (**Fig. 4G and Supplementary Fig. S4F**). To further substantiate this observation, we examined the infiltration of CD8 T cells within B16F10 tumors, which are highly immunosuppressive and refractory to immune infiltration (36). Interestingly, immunization with senB16F10 cells enhanced the infiltration of CD8 T cells into B16F10 tumors (**Fig. 4H**). Taken together, our observations suggest that immunization with viable senescent cancer cells can promote a tumor prophylactic and therapeutic CD8-dependent anti-tumor immune response.

### Senescent cancer cells hyper-stimulate reactive TILs from human patients

Finally, we validated our findings using a clinically relevant human cancer model. Therefore, we investigated whether the induction of senescence in patient-derived tumor cells enhances the activation of autologous tumor-infiltrating lymphocytes (TILs). To address this, TILs from different fragments of primary tumors were expanded *ex vivo* and tested for recognition of their autologous tumor cells. Based on these results, the different subpopulations of TILs were classified as tumor-reactive or non-reactive (**Fig. 5A**). This process was performed using therapy-naïve head and neck tumors from two different patients, VHIO-008 and VHIO-009. Upon treatment with anti-CD3-coated beads (OKT3), both tumor-reactive and non-reactive TILs showed robust upregulation of the activation marker 4-1BB (also known as CD137), indicating that both types of TILs were functional (**Fig. 5B and C**). As expected, VHIO-008 and VHIO-009 cancer cells activated their autologous reactive TILs but not their non-reactive TILs (**Fig. 5B and C**). Interestingly, *ex vivo* induction of senescence in these cancer cells with bleomycin (**Supplementary Fig. S5A**) led to an even stronger activation of reactive TILs (2- to 3-fold higher) compared to non-senescent cancer cells (**Fig. 5B and C**). Notably, senescent cancer cells did not stimulate non-reactive TILs, further indicating that TIL stimulation by senescent cells requires antigen recognition. These observations were validated in an antigen-specific setting. We used enriched populations of TILs from patient VHIO-008 specific to MAGEB2_p.E167Q_ and RPL14_p.H20Y_, two neoantigens previously identified by whole exome sequencing of the autologous tumor cell line. Neoantigen-specific TILs were o-cultured either with VHIO-008 parental or senescent tumor cells and 4-1BB expression was assessed. In this case, TIL activation with senescent VHIO-008 was more potent (2-6-fold higher) than that when using in non-senescent parental cells (**Fig. 5D**). Of note, senescence induction strongly upregulated MHC-I in VHIO-008 cells (**Supplementary Fig. S5C**). However, VHIO-009 cells had constitutively high levels of MHC-I, that were unaffected by the induction of senescence (**Supplementary Fig. S5D**). Together, these data indicate that the induction of senescence in human tumor cells potentiates antigen-dependent CD8 T cell activation.

**Figure 5:**
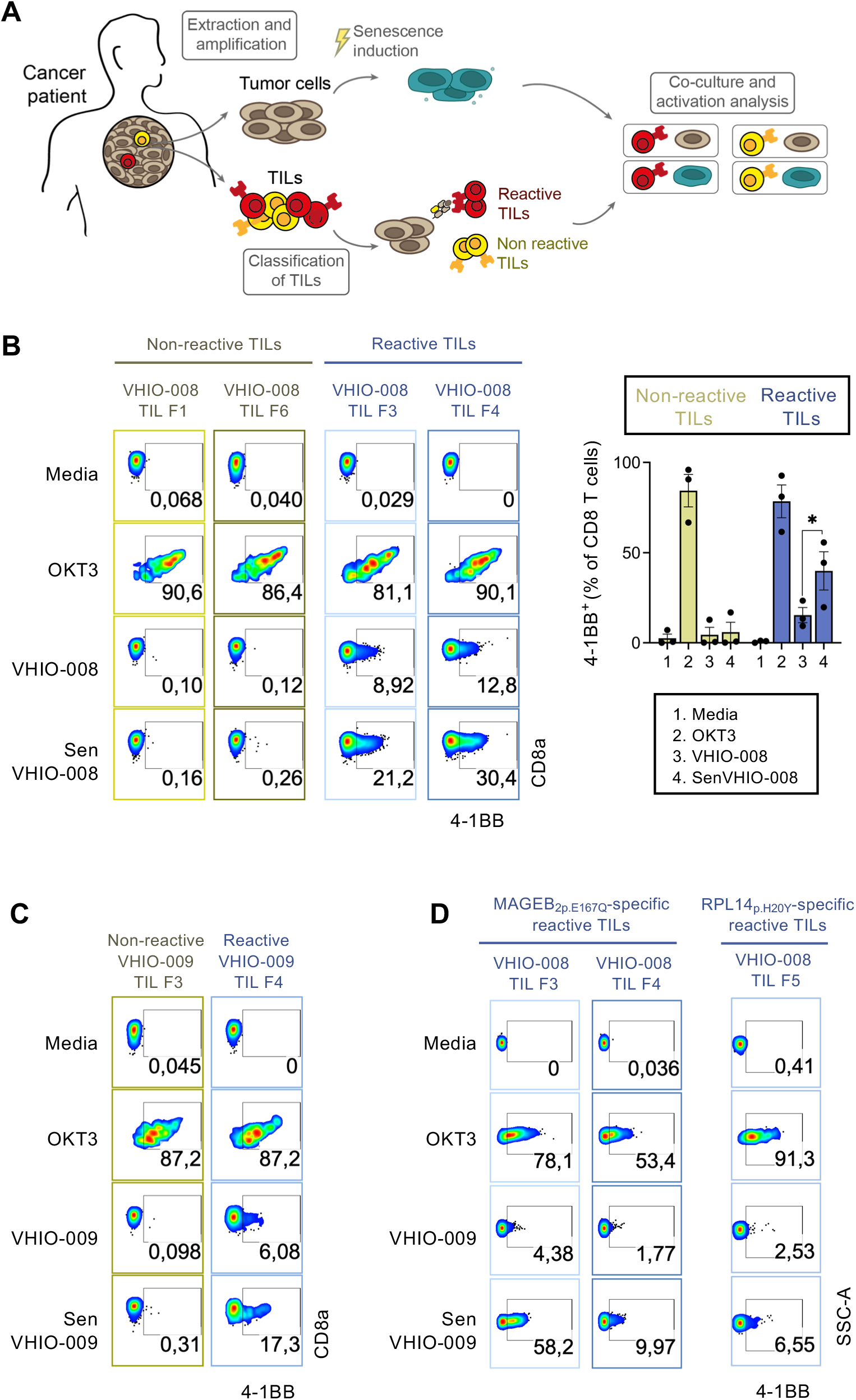
Cancer senescent cells hyper-stimulate autologous reactive TILs from human patients. A. Schematics of the procedure for isolating, amplifying, classifying and co-culturing patient derived tumor cells with autologous reactive and non-reactive TILs. B. Flow cytometry analysis of the activation marker 4-1BB in CD8 non-reactive (fragments F1 and F6) and reactive (fragments F3 and F4) autologous TILs from patient VHIO-008 after co-culture with RPMI media either alone or with anti-CD3 (OKT3), untreated VHIO-008 cells or bleomycin-treated senescent VHIO-008 cells (as indicated). The pseudocolor plots correspond to one representative experiment. Quantification of a total of 3 independent biological replicates with F6 and F3 TILs is shown in the right side. **p<0.01, unpaired Student’s t test C. Flow cytometry analysis of the activation marker 4-1BB in CD8 non-reactive (fragment F3) and reactive (fragment F4) autologous TILs from patient VHIO-009 after co-culture with RPMI media either alone or with anti-CD3 (OKT3), untreated VHIO-009 cells or bleomycin-treated senescent VHIO-009 cells (as indicated). Representative pseudocolor plots are shown. D. Flow cytometry analysis of the activation marker 4-1BB in CD8 reactive autologous TILs (reactive F3, F4 or F5 fragments) enriched to be reactive against to MAGEB2_p.E167Q_ and RPL14_p.H20Y_ (two neoantigens previously identified by whole exome sequencing of the autologous tumor cell line) after co-culture with RPMI media either alone or with anti-CD3 (OKT3), untreated VHIO-008 cells or bleomycin-treated senescent VHIO-008 cells (as indicated). Representative pseudocolor plots are shown.

## DISCUSSION

The ability of deleterious cells to drive adaptive immunity is strictly associated with two factors, adjuvanticity and antigenicity (33). Danger signals acting as adjuvants are essential to activate adaptive immune cells, and if absent, presentation of antigenic peptides to T cells drives peripheral tolerance rather than immune activation (30,31,34). Conversely, adjuvant signals in the absence of antigenic determinants drive inflammation but not adaptive immune responses. In this study, we show that senescent cells exhibit strong immunogenic potential, combining enhanced antigenicity and strong and sustained adjuvanticity, which together can drive CD8-dependent immune responses.

We compared a number of adjuvant-related features in parallel between senescent cells and cells undergoing immunogenic cell death (ICD), which constitute the current gold standard for eliciting cell-based anti-tumor immune responses (33,34). The levels of alarmins released into the extracellular milieu, particularly ATP and CALR, were similar between senescent and ICD cells, and this was reflected by the abundant recruitment of myeloid cells when injected subcutaneously. Notably, senescent cells outperformed ICD cells in the following two aspects. First, senescent cancer cells had a long persistence in the skin for up to 9 days post-injection, whereas ICD cells were essentially undetectable after the first day post-injection. Second, senescent cells were superior to ICD cells in transferring cellular material to dendritic cells, both from the cytosol and cytoplasmic membrane.

Regarding antigenicity, we initially focused on the expression of MHC-I, which we identified as upregulated through an unbiased approach *vis-a-vis* proteomic screening comparing senescent and non-senescent cells. Remarkably, senescent cells also upregulated the MHC-I-associated presentation machinery, including its master transcriptional regulator, *Nlrc5* (23). In agreement with previous observations (22), we also found that the interferon transcriptomic signature was elevated in senescent cells. Interferons are considered to be the main inducer of MHC-I presentation (21); therefore, they can explain our observations.

Interestingly, we show that immunization with non-cancer syngeneic senescent cells triggers CD8 T cell responses against both senescent and non-senescent cells. To pin-point senescence-specific responses, we wondered if we could identify senescence-associated MHC-I-peptides. The immunopeptidome presented by the MHC-I reflects a minority (at most 1%) of all the possible peptides from the proteome (36), and it is also highly dynamic depending on the cellular context and, particularly, under conditions of stress (34,38). We speculated that senescent cells, even if derived from non-cancer cells, could present senescence-associated peptides. Indeed, by isolating and sequencing the immunopeptidome of senescent cells, we demonstrated the presence of senescence-associated peptides that were not detected in their parental non-senescent cells or in the Mouse Immunopeptidome Atlas (which includes >30,000 peptides normally present in a large collection of murine tissues and a set of cancer cell lines) (29). Moreover, CD8 T cells from mice immunized with senescent cells were activated *ex vivo* when exposed to senescence-specific peptides. This was not the case when mice were immunized with parental non-senescent cells. These observations indicate that senescent cells have the capacity to elicit CD8 T-cell responses partly evoked by senescence-associated peptide antigens.

Having demonstrated that non-cancer senescent cells are immunogenic, we extended this concept to cancer cells. We showed that senescent cancer cells are superior to ICD cancer cells in triggering protection against a subsequent challenge with cancer cells. As expected, anti-tumor protection involved a classical adaptive immune response mediated by antigen-presenting cells (CD11b^+^) and CD8 T cells. We also compared the immunogenic potency of live *versus* dying senescent cancer cells, and observed that live senescent cells were more efficient. This finding further reinforces the idea that senescent cells play an active role in triggering an efficient immune response. We also applied this strategy in a therapeutic setting by immunizing tumor-bearing mice. In this case, immunization significantly delayed B16F10 melanoma growth, which is considered a highly immunosuppressive tumor model (36). This delay in tumor growth was accompanied by a significant infiltration of CD8 T cells within the tumors.

The above concepts were corroborated in a human setting based on autologous TILs and tumor cells from primary isolates from human cancer patients. We found that co-culture of TILs with senescent cancer cells evokes stronger antigen-dependent activation than co-cultured with their non-senescent parental cells. We also demonstrated this in the case of TILs that recognize a single cancer-derived mutated antigen.

Our findings demonstrate that senescent cells are strongly immunogenic owing to the combination of several features: their long-term persistence *in vivo*, their release of adjuvant factors, their efficient transfer of cellular material to APCs, and their altered immunopeptidome. These results suggest the possibility of improving anti-cancer vaccination strategies. Furthermore, unleashing adaptive immune responses or engineered TCR-T cells against senescent cells may have therapeutic benefits for diseases in which senescent cells play a pathological role.

## Methods

### Cell culture

SKMEL-103 (human melanoma), IMR-90 (human lung fibroblasts), and B16-F10 (mouse melanoma) cells were obtained from the ATCC). Panc02 (mouse pancreatic adenocarcinoma) cells were kindly provided by Dr. Azad (Oxford University). OVA-expressing B16-F10 cells were kindly provided by the Kroemer Lab (CRC, Paris). Luciferase-expressing B16-F10 cells were kindly provided by the Soengas Lab (CNIO, Madrid). Primary mouse embryonic fibroblasts (MEF) were obtained from C57BL/6J embryos at E13.5 as previously described (39). Panc02 cells and all primary immune cells were maintained in standard RPMI medium. All the other cell lines were maintained in standard DMEM. All media were supplemented with 10% heat inactivated Fetal Bovine serum (FBS) (Gibco) and 1% antibiotics (penicillin/streptomycin 100 U/mL; Gibco). Cells were maintained in a humidified incubator at 37 °C and 5% CO_2_. Cells were routinely tested for mycoplasma contamination using standard PCR, and only negative cells were used.

For senescence induction, cells were treated with doxorubicin (200 nM) (Sigma #D1515) for 48 h (after that time, fresh media was added), palbociclib (1 μM) (PD033299, #S1116 Absource Diagnostic) or nutlin-3A (5 μM) (Sigma, #SML0580), as indicated. 7-10 days after the treatment, senescent cells were collected and used for experiments. Late passage senescence was triggered by repeated subculture of non-transformed primary cells. ICD was induced by treatment with a high dose of doxorubicin (5 μM) for 18h; after that, cells were corrected and used for experiments. Senescent cell death was induced by treatment with the senolytic agent navitoclax (10 μM) for 18h; after that, cells were corrected and used for experiments.

### Senescence-associated β-galactosidase assay

Cells were washed with PBS and fixed with 0.2% glutaraldehyde. After washing with PBS, cells were incubated overnight at 37 °C with a staining solution containing 1 mg/mL X-Gal (Melford BioLaboratories, MB1001) prepared in dimethylformamide (DMF, Sigma, D4551) at pH 6. Cells were then washed in PBS and visualized using a Nikon Eclipse TS2 bright-field microscope.

### Plasma membrane proteomic screening

Up to 5×10^6^ cells per condition were collected in cold PBS by scraping, and pelleted by centrifugation. Plasma membrane proteins were extracted using a plasma membrane protein extraction kit (Abcam, #ab65400), following the manufacturer’s instructions.

Proteins were dissolved in UT buffer (8 M Urea, 2 M thiourea, 100 mM Tris-HCl pH=8.0) and digested by means of the standard FASP protocol. Briefly, proteins were reduced (15 mM TCEP, 30 min, RT), alkylated (50 mM CAA, 20 min in the dark, RT) and sequentially digested with Lys-C (Wako) (protein:enzyme ratio 1:50, o/n at RT) and trypsin (Promega) (protein:enzyme ratio 1:100, 6 h at 37 °C). Resulting peptides were desalted using Sep-Pak C18 cartridges (Waters). LC-MS/MS was done by coupling an UltiMate 3000 RSLCnano LC system to either a Q Exactive HF or Q Exactive HF-X-mass spectrometer (Thermo Fisher Scientific). In both cases, peptides were loaded into a trap column (Acclaim™ PepMap™ 100 C18 LC Columns 5 µm, 20 mm length) for 3 min at a flow rate of 10 µl/min in 0.1% FA. Then, peptides were transferred to an EASY-Spray PepMap RSLC C18 column (Thermo) (2 µm, 75 µm x 50 cm) operated at 45 °C and separated using a 90 minute effective gradient (buffer A: 0.1% FA; buffer B: 100% ACN, 0.1% FA) at a flow rate of 250 nL/min. The gradient used was: from 4% to 6% of buffer B in 2.5 min, from 6% to 25% B in 72.5 min, from 25% to 42.5% B in 14 min plus 6 additional minutes at 98% B. Peptides were sprayed at 1.8 kV into the mass spectrometer via the EASY-Spray source and the capillary temperature was set to 300 °C. The Q Exactive HF was operated in a data-dependent mode, with an automatic switch between MS and MS/MS scans using a top 15 method. (Intensity threshold ≥ 6.7e4, dynamic exclusion of 26.25 sec and excluding charges +1 and > +6). MS spectra were acquired from 350 to 1400 m/z with a resolution of 60,000 FWHM (200 m/z). Ion peptides were isolated using a 2.0 Th window and fragmented using higher-energy collisional dissociation (HCD) with a normalized collision energy of 27. MS/MS spectra resolution was set to 15,000 or 30,000 (200 m/z). The ion target values were 3e6 for MS (maximum IT of 25 ms) and 1e5 for MS/MS (maximum IT of 15 or 45 msec). The Q Exactive HF-X was operated in a data-dependent mode, with an automatic switch between MS and MS/MS scans using a top 12 method. (Intensity threshold ≥ 3.6e5, dynamic exclusion of 34 sec and excluding charges +1 and > +6). MS spectra were acquired from 350 to 1400 m/z with a resolution of 60,000 FWHM (200 m/z). Ion peptides were isolated using a 1.6 Th window and fragmented using higher-energy collisional dissociation (HCD) with a normalized collision energy of 27. MS/MS spectra resolution was set to 15,000 (200 m/z). The ion target values were 3e6 for MS (maximum IT of 25 ms) and 1×10^5^ for MS/MS (maximum IT of 22 msec). Raw files were processed with MaxQuant using the standard settings against either a human protein database (UniProtKB/Swiss-Prot, 20373 sequences) or a mouse database (UniProtKB/TrEMBL, 53449 sequences). Carbamidomethylation of cysteines was set as a fixed modification whereas oxidation of methionines and protein N-term acetylation were set as variable modifications. Minimal peptide length was set to 7 amino acids and a maximum of two tryptic missed-cleavages were allowed. Results were filtered at 0.01 FDR (peptide and protein level). Afterwards, the “proteinGroups.txt” file was loaded in Prostar (Wieczorek et al, Bioinformatics 2017) using the LFQ intensity values for further statistical analysis. Briefly, proteins with less than 75% valid values in at least one experimental condition were filtered out. When needed, a global normalization of log2-transformed intensities across samples was performed using the LOESS function. Missing values were imputed using the algorithms SLSA (40) for partially observed values and DetQuantile for values missing on an entire condition. Differential analysis was performed using the empirical Bayes statistics Limma. Proteins with a p-value < 0.05 and a log_2_ ratio >0.58 (1.5 in non-log scale) were defined as upregulated. The FDR was estimated to be below 5%. Upregulated proteins in senescent cells following the indicated criteria in four or more conditions of senescence were selected, and GO analysis was performed using the GOrilla Gene Ontology tool (41).

### Flow cytometry analysis

For analysis of cultured cell lines, single cells were digested into single cells by trypsinization (0.25% trypsin-EDTA, Invitrogen). Non-adherent primary immune cells maintained in suspension were collected from the culture by pipetting. Adherent primary immune cells (BMDC) were collected by cell scraping. Blood was collected in EDTA-coated tubes (16.444, Microvette®) to assess immune cell depletion and incubated with RBC lysis buffer (42031, BioLegend) for 5 min at room temperature (RT). Cells were resuspended in fluorescence-activated cell sorting (FACS) buffer (5 mM EDTA and 0.5% BSA in PBS). Cell viability was assessed using Live/Dead Fixable Yellow dye (Invitrogen) following the manufacturer’s instructions or with 0.1 µM DAPI (Molecular Probes, D1306). Dead cells were excluded from the analysis. Mouse cells were incubated with mouse BD Fc BlockTM containing purified anti-mouse CD16/CD32 mAb 2.4G2 at 1:400 (553142, BD BioSciences) for 10 min at 4 °C. After washing, cells were stained with the appropriate antibody (**Supplementary Table S4**) for 40 min at 4 °C. When necessary, a fluorophore-conjugated secondary antibody (#A11029; Invitrogen) was used for 40 min at 4 °C. Cell suspensions were run on a Gallios Beckman Coulter flow cytometer (BD Biosciences). Autofluorescence signals from the unstained samples were obtained and subtracted from each sample in all experiments. Data were analyzed using FlowJo v10 software.

### RNA extraction, RNA-seq library preparation and sequencing

Total RNA was extracted from untreated proliferating and doxorubicin-treated senescent MEF using the RNeasy® Mini Kit (Qiagen), following the manufacturer’s instructions. Library preparation and quality control were performed at the Genomics Facility of the IRB Barcelona. The total RNA concentration was quantified using the Qubit RNA HS Assay kit (Invitrogen), and RNA integrity was assessed using the Bioanalyzer 2100 RNA Nano assay (Agilent). RNA-Seq libraries were prepared at the IRB Barcelona Functional Genomics Core Facility. Briefly, mRNA was isolated from 1.1 μg total RNA using the NEBNext Poly(A) mRNA Magnetic Isolation Module (New England Biolabs). Isolated mRNA was used to generate dual-indexed cDNA libraries using the NEBNext Ultra II Directional RNA Library Prep Kit for Illumina (New England Biolabs). Eight cycles of PCR amplification were applied to all the libraries. The final libraries were quantified using the Qubit dsDNA HS assay (Invitrogen), and the quality was controlled using the Bioanalyzer 2100 DNA HS assay (Agilent). An equimolar pool was prepared using the 13 libraries and submitted for sequencing at the National Centre for Genomic Analysis (CRG-CNAG). Final quality control by qPCR was performed by the sequencing provider before paired-end 150 nt sequencing on a NovaSeq6000 S4 (Illumina). The sequencing results exceeded 312Gbp, with a minimum of 53.97 million paired-end reads sequenced for each sample. Adapters were trimmed from the initial sequences using Cutadapt (42), version 1.18. Trimmed paired-end reads were then aligned to the mm10 version of the mouse genome using STAR (43) version 2.3.0e under default parameters. SAM files were sorted and indexed using Sambamba (44) version 0.5.9. R (45) package Casper (46), version 2.16.1, was used to quantify the intensities at the transcript and gene levels. The ENSEMBL database was used for the transcript annotation. Differential expression between conditions senMEF and control MEF was performed using the limma R package (47), version 3.38.2, on the gene level intensities using the replicate Id as an adjusting covariate to account for paired samples. Genes were annotated to hallmark terms (48) using the R package org.Hs.eg.db (49). Gene set analysis was then performed to filter out low-expression genes (genes with less than an average count of 5 reads). The rotation-based approach for enrichment (50) implemented in the R package limma was used to represent null distribution. The maxmean enrichment statistic proposed in (51) under re-standardization was considered for competitive testing.

### mRNA levels analysis

Total RNA was extracted from cell samples using TRIzol reagent (Invitrogen) according to the manufacturer’s instructions. Up to 1 µg of total RNA was reverse transcribed into cDNA using the iScript cDNA Synthesis Kit (Bio-Rad) or gDNA Clear iScript cDNA Synthesis Kit for patient-derived tumor cells (Bio-Rad) for RT-qPCR. Quantitative real-time PCR was performed using GoTaq PCR Master Mix (Promega) or PowerUp SYBR Green Master Mix (Thermo Fisher Scientific) for patient-derived tumor cells in a QuantStudio 6 Flex thermicycler (Applied Biosystems) 7900HT Fast Real-Time PCR System (Applied Biosystems). The average expression of both endogenous *Actb* and *Gapdh* in mouse cells (*ACTB* and *GAPDH* in human cells) genes served as endogenous normalization controls. for patient-derived tumor cells, and expression of the endogenous *GAPDH* gene served as an endogenous normalization control. Primers used in this study are listed in **Supplementary Table S4**.

### Immunopeptidome

Up to 10^8^ cells per condition were collected by trypsinization and pelleted by centrifugation. Cells were washed with PBS, thawed at −80 °C, and shipped to the CHU Sainte-Justine Research Center (Montreal) for immunopeptidomics analysis using mass spectrometry (52).

Materials and reagents: Anti-Mouse H2 (M1/42.3.9.8, #BE0077) from BioXcell, Polyprep chromatography column (#7311553), and combined inhibitor EDTA-free (#A32961) from Bio-Rad and solid phase extraction disk ultramicrospin column C18 (#SEMSS18V, 5-200 µL) from the nest group. 1.5 mL and 2.0 mL microcentrifuge tubes (Protein LoBind Eppendorf #022431081 and #02243100), low retention tips Eppendorf (10 µL #2717349, 20 µL #2717351, 200 µL #2717352), acetonitrile (#A9964), trifluoroacetic acid (TFA, #AA446305Y), formic acid (#AC147930010), chaps (#22020110GM), PBS (Buph, phosphate buffer saline packs, #28372), CNBr-activated sepharose 4 B (#45000066), and ammonium bicarbonate (#A643-500) were purchased from Fisher.

To isolate MHC class I peptides, a frozen pellet of 10^8^ cells was thawed by warming the bottom of the tube with palm. The cell pellet was resuspended in 500 µL of PBS by pipetting up and down until homogenization. The volume of the cell pellet suspension was measured and transferred into a new 2 mL microcentrifuge tube. An equivalent volume of cell lysis buffer (1% CHAPS in PBS containing protease inhibitors, 1 pellet/10 mL) was added to the cell suspension (final concentration of the lysis buffer of 0.5% Chaps), followed by an incubation for 60 min at 4 °C using a rotator device (10RPM) and centrifugation at 18,000 × g for 20 min at 4 °C. The cell lysis supernatant containing the MHC-peptide complexes was transferred to a new 2 mL microcentrifuge tube and kept on ice until use for immunopurification.

Bead coupling and immunopurification of MHC class I peptides were performed as previously described (53). Briefly, 80 mg of sepharose CNBr activated beads was coupled with 2 mg of anti-mouse H2 antibody. Sepharose antibody-coupled beads were incubated with the cell lysate supernatant in a 2 mL Low binding microcentrifuge tube overnight at 4 °C with rotation (22 RPM). The next day, a Bio-Rad column was installed on a rack and pre-rinsed with 10 mL of buffer A (150 mM NaCl and 20 mM Tris-HCl –HCl pH 8). The bead-lysate mixture was transferred into a Bio-Rad column, and the bottom cap was removed to discard the unbound cell lysate. The beads retained in the Bio-Rad column were washed sequentially with 10 mL of buffer A (150 mM NaCl and 20 mM Tris–HCl pH 8), 10 mL of buffer B (400mM NaCl and 20 mM Tris–HCl pH 8), 10 mL of buffer A, and 10 mL of buffer C (20 mM Tris–HCl pH 8.). MHC-peptide complexes were eluted from the beads by adding 300 µL of 1% TFA, pipetting up and down 4-5 times and collecting the flow-through. This step was repeated once, and the flowthroughs were collected and combined in a new 2 mL tube.

MHC class I peptides were desalted and eluted on a C18 column. First, the C18 column was pre-conditioned with 200 µL of 1) methanol, 2) 80% acetonitrile/0.1% TFA and 3) 0.1%TFA and spun at 1545 g in a fixed rotor to collect and discard the flowthroughs. Then, The MHC-peptide complexes previously collected in 600 µL of 1% TFA were loaded (3 × 200 µL) into the pre-conditioned C18 column, spun and flowthroughs were discarded. The final wash was performed with 200 µL of 0.1% TFA and spun again. Finally, the C18 column was transferred onto a 2 mL Eppendorf tube, and MHC class I peptides were eluted with 3 × 200 µL of 28%ACN 0.1%TFA. The flow-through containing the eluted peptides was stored at −20 °C for MS analysis. Prior to LC-MS/MS analysis, the purified MHC class I peptides were evaporated to dryness using a vacuum concentrator with presets of temperature 45 °C for 2 h, vacuum level:100 mTorr, and vacuum ramp:5.

Vacuum samples were resuspended in 52 µL of 4% formic acid (FA), and each biological replicate was divided into three technical replicates of 16µl each. Each replicate was loaded and separated on a homemade reversed-phase column (150 μm i.d. × 250 mm length, Jupiter 3 µm C18 300 Å) with a gradient from 10 to 30% ACN-0.2% FA and a 600-nl/min flow rate on an Easy nLC-1000 connected to an Orbitrap Eclipse (Thermo Fisher Scientific). Each full MS spectrum was acquired at a resolution of 240000, an AGC of 4E5, and an injection time of 50ms, followed by tandem-MS (MS-MS) spectra acquisition on the most abundant multiply charged precursor ions for a maximum of 3 s. Tandem-MS experiments were performed using higher energy collisional dissociation (HCD) at a collision energy of 34%, a resolution of 30000, an AGC of 1.5E4, and an injection time of 300ms. Data files were processed using PEAKS X software (Peaks Pro V10.6, Bioinformatics Solutions, Waterloo, ON, USA) using the mouse database UniProtKB/Swiss-Prot (2019_09). ‘Unspecified enzyme digestion’ was selected for the enzyme parameter and mass tolerances on precursor and fragment ions were 10ppm and 0.01 Da, respectively. All other search parameters were the default values. Final peptide lists were filtered using ALC of 80% and with a false discovery rate (FDR) of 1% using the Peaks software. Data were visualized using MhcVizPipe software (54). Cross-analysis to identify unique peptides was performed using Venn diagrams (55) and the previously described H-2K^b^/D^b^ immunopeptidome atlas generated from mouse primary tissues (29).

Only captured peptides between 8- to 12-mers (both included) and those predicted to be weak or strong binders to either H-2K^b^ or H-2K^d^ (NetMHCPan v4.0) were considered for the analysis (*bona fide* peptides). We combined all *bona fide* peptides detected as senescent MEF and removed all peptides detected in non-senescent cells. Peptides present in other tissues and cancer cell lines of C57BL/6J mice were obtained from the Mouse Immunopeptidome Atlas (29) and subtracted to filter our list of candidate peptides. For the list of candidate peptides, we explored the senescent MEF *versus* MEF differences in mRNA for their predicted genes (under reverse translation) using RNA-seq data, giving extra priority to those peptides whose underlying genes were upregulated in senescent cells.

### Mice

All mice were maintained at the animal facility of the Scientific Parc of Barcelona (PCB) in strict accordance with Spanish and European Union regulations. Animal experiments were approved by the Animal Care and Use Ethical Committee of the PCB and Catalan Government. Mice were maintained under specific pathogen-free conditions. Food and water were provided *ad libitum*. All *in vivo* experiments were performed using female C57BL/6J mice of 8-16 weeks of age, that were randomly allocated in the different groups of study. All mice were purchased from Charles Rives, France.

### Mouse immunizations

For immunization with non-cancer cells, 8-12 weeks C57BL/6J female mice were subcutaneously injected with 10^6^ untreated or senescent MEF resuspended in 100 μL PBS together with 25 μg of CpG immune adjuvant (synthetic oligodeoxynucleotide [ODT] containing unmethylated CpG motifs; ODT1825 Vaccigrade, ac-1826-1) on days 0 and 7 on the left and right dorsal flanks, respectively. As negative controls, animals received only the vehicle plus adjuvant, and as a positive control, animals were immunized with 100 μg of ovalbumin (A5503, Merck Life Sciences) in a total volume of 100 μL PBS. One week after the last immunization, the animals were euthanized by CO_2_, and their spleens were used for the analysis of immune responses.

To analyze the adjuvant properties of cancer cells, 8-12 weeks C57BL/6J female mice were subcutaneously injected with vehicle (PBS), 10^6^ ICD-cells, or 2×10^5^ senescent cells resuspended in 100 μL PBS on day 0 on the left flank. On day 7, the subcutaneous inoculation site was dissected and subjected to histological analysis.

For prophylactic immunization with cancer cells, 8-12 weeks C57BL/6J female mice were subcutaneously injected with 10^6^ ICD cells or 2×10^5^ senescent cells (either live or dead, as indicated) resuspended in 100 μL PBS on day 0 on the left flank. On day 7, the animals were re-challenged with 3×10^5^ live cancer cells in their right flank. Tumor appearance and growth were monitored afterwards.

For depletion of immune populations, animals received 100 μg of blocking antibodies (anti-CD4 (clone GK1.5, BioxCell), anti-CD8 (clone 2.43, BioxCell), anti-CD11b (clone M1/70, BioxCell)) or isotype control IgG2b (clone LTF-2, BioxCell) via intraperitoneal injection of a total volume of 100 μL on days -1, 3, and 8 of the immunization experiments. For therapeutic vaccination, the animals were injected with 5×10^5^ live cancer cells in the right flank. When tumors were visible and palpable (from 40mm^3^), animals were immunized subcutaneously with 10^6^ ICD cells or 2×10^5^ senescent cells (either live or dead, as indicated) resuspended in 100 μL PBS on the left flank. Tumors were measured using a caliper, and their volumes (v) were calculated using the formula v=(lxw^2^)/2 (where l is tumor length and w is tumor width). The measurements are shown in mm^3^. When tumors reached a size of 1000 mm^3^ or became ulcerated, mice were euthanized by CO_2_ and samples were collected for further analysis.

### Histological analysis

For formalin-fixed paraffin-embedded (FFPE) samples, tissues were fixed overnight at 4 °C with neutral buffered formalin (HT501128-4L, Sigma-Aldrich). Paraffin-embedded tissue sections (2-3 μm) were air-dried and dried overnight at 60 °C.

For hematoxylin-eosin (H/E) staining, paraffin-embedded tissue sections were dewaxed and stained according to the H/E standard protocol using a CoverStainer (Dako, Agilent).

Immunohistochemistry for CD45 (30-F11) (14-0451-82, ThermoFisher) at 1:100 for 60 min was performed using a Ventana Discovery XT, and for CD3 (IR503, Dako-Agilent) at 1:10 and 120 min, CD11b [EPR1344] (ab133357, Abcam) at 1:6000 for 120 min, and CD8 alpha [EPR20305] (ab209775, Abcam) at 1:1000, 120 min with the Leica BOND RX. For CD45, antigen retrieval was performed using Cell Conditioning 1 (CC1) buffer (950-124, Roche) followed by rabbit anti-Rat (AI-4001, Vector) at 1:500 for 32 min and OmniMap™ anti-Rb HRP (760-4311, Roche). Blocking was performed using casein (760-219, Roche). Antigen–antibody complexes were revealed using the Discovery Purple Kit (760-229, Roche). For CD3, CD11b, and CD8 alpha antigen retrieval was performed with BOND Epitope Retrieval 2 – ER2 (AR9640, Leica) for 40, 20, and 20 min, respectively, followed by Bond Polymer Refine Red Detection (DS9390, Leica) without the post-primary for 30 min. The sections were mounted with mounting medium and toluene-free (CS705, Agilent) using a Dako CoverStainer. The specificity of staining was confirmed by staining with rat IgG (ref:6-001-F, R&D Systems, Biotechne) or rabbit IgG (ab27478, Abcam) as an isotype control. Brightfield images were acquired using a NanoZoomer-2.0 HT C9600 digital scanner (Hamamatsu) equipped with a 20X objective. All images were visualized with a gamma correction set at 1.8 in the image control panel of the NDP.view 2 U12388-01 software (Hamamatsu, Photonics, France). Brightfield immunohistochemistry images were blindly quantified using QuPath Software o.1.2 (56).

### ELISpot assay

To test immune responses in animals immunized with non-cancer senescent cells (MEF), splenocytes from naïve (injected with vehicle plus adjuvant) and immunized animals (injected with untreated, senescent cells, or ovalbumin plus adjuvant) were collected 7 to 10 days after the last immunization. Briefly, spleens were harvested, and RBC were lysed using RBC lysis buffer (#40301, Palex Medical SA) and seeded as the effector cell population at 1.5×10^5^ cells per well into mouse IFN-γ ELISpot plates (15607416, Fisher Scientific). Target cells (control or senescent cells) were added and co-cultured at a 1:10 ratio (target:splenocytes). Peptides were added at concentrations of 400 nM (when added alone) and 200 nM (when added in pools) to the cultures. The OVA-derived peptide SIINFEKL (S795, Sigma) was used as the positive control (400 nM). Peptides obtained from immunopeptidomes were synthesized using Pepscan. Stimulation was maintained for 20 h. Afterwards, plates were developed according to the manufacturer’s protocol. They were scanned and quantified using Elispot7.0iSpot software in an ImmunoSpot Plate Reader.

### Measurement of ATP levels

An equal volume of fresh medium was added to 10^6^ cells per condition. 24 h after, the culture medium was collected, centrifuged, and filtered through a 0.2 μm filter to eliminate cell debris. The medium was analyzed using CellTiterGlo® (G7571, Promega), according to the manufacturer’s protocol. The medium alone was used for background control of luminescence, and the signal was subtracted from the samples.

### Measurement of CALR secretion

10^6^ cells per condition were washed three times with PBS, and FBS-free culture medium was added. 24h later, the conditioned media (CM) was collected and concentrated using AMICON Ultra-15 tubes (UFC900324, Merck Life Sciences) by centrifugation for 1 h at 4 °C. Up to 20 μL of concentrated CM per sample was loaded per lane and hybridized using antibodies against CALR (9707, Cell Signalling). Secondary fluorescent reagents (Goat anti-mouse or anti-rabbit IRDye 680RD and 800CW, LI-COR) were used according to the manufacturer’s instructions.

### BMDC generation

Bone marrow was harvested from the femurs and tibias of the C57BL/6 mice. The bone marrow was flushed and centrifuged at 350 × g for 5 min. Blood cells were lysed by incubation with RBC lysis buffer for 5 min at room temperature. Cells were filtered sequentially through 100 μm and 70 μm filters to remove aggregates. Cells were resuspended and cultured on 10 cm^2^ bacterial Petri dishes in RPMI with 20 ng/mL murine GM-CSF (315-03, PeproTech). On day 3, fresh medium containing fresh GM-CSF was added, and the medium was replaced on day 6. BMDC were harvested on day 7 by gentle pipetting and used for the assays.

### Assessment of BMDC antigen capture

Untreated, ICD, and senescent cancer cells were stained using the fluorescent cytosolic-dye CellTrace CFSE (Thermofisher) or the fluorescent membrane-dye wheat germ agglutinin WGA-Alexa Fluor 647 (Thermofisher). Fluorescently labeled cancer cells were then co-cultured with BMDCs in a 1:1 ratio for 18h, and the BMDCs were analyzed by flow cytometry for acquisition of CFSE or WGA-Alexa-647.

### Patient characteristics and patient-derived tumor samples

Tumor specimens were obtained from two patients with metastatic head and neck cancer: VHIO-008 had metastatic hypopharyngeal carcinoma and VHIO-009 had metastatic oropharyngeal carcinoma. Both patients were refractory to standard lines of therapy prior to sample procurement. Informed consent was obtained from all subjects. Tumor biopsies were cut into small 2-4 mm fragments to expand independent TIL lines in the presence of 6000 IU/mL IL-2. Tumor cell lines were established by culturing one tumor fragment in RPMI 1640 supplemented with 20% human AB serum (Biowest), 100 U/mL penicillin, 100 μg/mL streptomycin, 25 mM HEPES (Thermo Fisher Scientific), 10 μg/mL gentamicin and 1.25 μg/mL amphotericin B at 37 °C and 5% CO_2_. After one month, adherent and non-adherent cells were cultured in RPMI 1640 supplemented with 20% FBS (Hyclone), 100 U/mL penicillin, 100 μg/mL streptomycin, 25 mM HEPES (Thermo Fisher Scientific), 10 μg/mL gentamicin (Lonza) and 1.25 μg/mL amphotericin B (Gibco). Occasionally, we used differential trypsinization to enrich for epithelial cells. Media was changed once a month until tumor cell line was established. Tumor cell lines were authenticated through whole exome sequencing.

Once established, VHIO-008 and VHIO-009 tumor cell lines were cultured in RPMI with GlutaMAX supplemented with 10% of fetal bovine serum (FBS) and 1% of Penicillin-Streptomycin (P/S). All cells were incubated at 37 °C, 5% CO_2_, in a humidified incubator. For senescence induction, cells were treated with 6 mU of bleomycin (Sigma-Aldrich, B8416) for 48 h. Then, cells were wash out twice with PBS and fresh media was added. After 5 days, cells were collected after trypsinization (Trypsin 0.05%) and further experiments were performed.

### Assessment of autologous tumor recognition by up-regulation of 4-1BB

To evaluate tumor recognition by TILs, autologous tumor cells were used as tumor targets in co-culture assays. 2×10^4^ *ex vivo* expanded TILs or TILs previously enriched for recognition of a specific neoantigen (i.e. MAGEB_2p.E167Q_ and RPL14_p.H20Y_) identified as previously described (57), were co-cultured with 5×10^4^ autologous tumor cells. After 20 h, T cell recognition was assessed by measuring the upregulation of the activation marker 4-1BB on the surface of T cells by flow cytometry. Briefly, co-cultured cells were pelleted, re-suspended in staining buffer, and incubated with the following antibodies for 30 minutes at 4°C: CD3 APC-H7 (1:20), CD8 Pe-Cy7 (1:60), 4-1BB APC (1:20). Cells were washed, resuspended in staining buffer containing propidium iodide (PI) and acquired on a BD LSRFortessa™. T cell reactivities were considered positive when the frequency of 4-1BB by flow cytometry was higher than the control. A positive control was included by stimulating the T cells with anti-CD3 antibody OKT3 (BioLegend).

### Statistical analysis

Data were analyzed using GraphPad Prism v.9.3.0 software and are represented as mean ± SEM of independent biological replicates. Statistical analyses were performed as described in the figures. Differences were considered significant based on P values (^#^p<0.1, *p<0.05, **p<0.01, ***p<0.001).

### Data availability

Data Access Statement: Research data supporting this publication are deposited in public repositories. The RNA sequencing data are available in Gene Expression Omnibus (GEO) located at https://www.ncbi.nlm.nih.gov/geo/query/acc.cgi?acc=GSE202032 with accession number GSE202032. The proteomic screen data is available at ProteomeXchange located at http://proteomecentral.proteomexchange.org/cgi/GetDataset. The immunopeptidomic data is available at Pride located at https://www.ebi.ac.uk/pride/archive/. Derived data supporting the findings of this study are available in supplementary tables as indicated.

## Supporting information

Proteomic screen of plasma membrane-enriched proteins in senescent cells

GO terms enrichment analysis of membrane proteins found upregulated in 4 or more senescence conditions

List of peptides presented exclusively on senescent cells MHC-I

## Author’s Contributions

**I. Marín**: conceptualization, data curation, software, formal analysis, validation, investigation, methodology, visualization, writing-original draft, and writing-review and editing. **O. Boix**: investigation and methodology for experimentation with patient-derived samples. **A. García**: investigation and methodology for experimentation with patient-derived samples. **I. Sirois**: investigation and methodology for immunopeptidomics. **A. Caballe**: software, formal analysis, and methodology for RNA sequencing and immunopeptidome data. **E. Zarzuela** software, formal analysis, and methodology for proteomic screen. **I. Ruano**: formal analysis and methodology of histological staining. **C. Stephan-Otto Attolini**: supervision for RNA sequencing and immunopeptidome data analysis. **N. Prats**: supervision, formal analysis, and methodology of histological staining. **J. A. López-Domínguez**: investigation for RNA-seq of Panc02 cells. **M. Kovatcheva**: investigation for RNA-seq of SKMEL-103 cells. **E. Garralda**: resources for experimentation with patient-derived samples. **J. Muñoz**: software, formal analysis, supervision, and methodology for proteomic screen. **E. Caron**: supervision and methodolgy of immunopeptidomics. **M. Abad**: conceptualization, resources, and supervision for experimentation with patient-derived samples. **A. Gros**: conceptualization, resources, and supervision for experimentation with patient-derived samples. **F. Pietrocola**: conceptualization, data curation, formal analysis, supervision, writing-original draft, and writing-review and editing. **M. Serrano**: conceptualization, resources, data curation, formal analysis, supervision, funding acquisition, writing-original draft, project administration, and writing-review and editing. All the authors discussed the results and commented on the manuscript.

## Acknowledgements

We are grateful to Maria Isabel Muñoz for assistance with the animal protocols, to the IRB core facilities (Functional Genomics, Biostatistics/Bioinformatics and Histopathology), and PCB (Animal House) for general research support. I.M. was funded by the Spanish government (FPI scholarship). J.A.L.-D. and M.K. were supported by a fellowship from the Spanish Association for Cancer (AECC). This work in the laboratory of E.C. was funded by the Fonds de recherche du Québec – Santé (FRQS), the Cole Foundation, CHU Sainte-Justine and the Charles-Bruneau Foundations, Canada Foundation for Innovation, the National Sciences and Engineering Research Council (NSERC) (#RGPIN-2020-05232), and the Canadian Institutes of Health Research (CIHR) (#174924). The work in the laboratory of F.P. is supported by a Karolinska Institute Starting Grant, Starting Grant from the Swedish Research Council (2019_02050_3), and by grants from the Harald Jeanssons Foundation, the Loo and Hans Osterman Foundation, and Cancerfonder (21 1637 Pj). Work in the laboratory of M.S. was funded by the IRB and “laCaixa” Foundation, and by grants from the Spanish Ministry of Science co-funded by the European Regional Development Fund (ERDF) (SAF2017-82613-R), European Research Council (ERC-2014-AdG/669622), and Secretaria d’Universitats i Recerca del Departament d’Empresa i Coneixement of Catalonia (Grup de Recerca consolidat 2017 SGR 282).

---

**Supplementary Figure S1:**
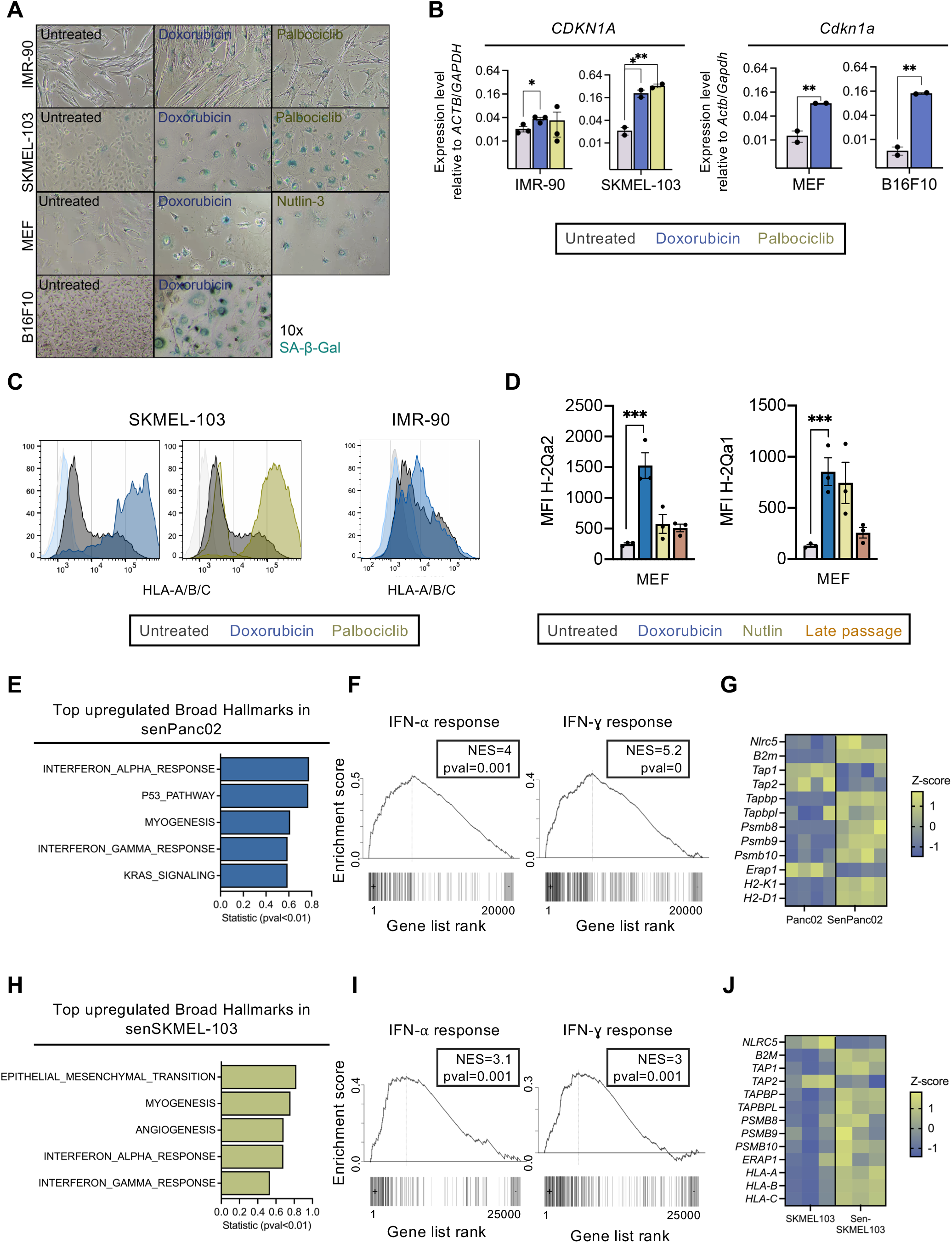
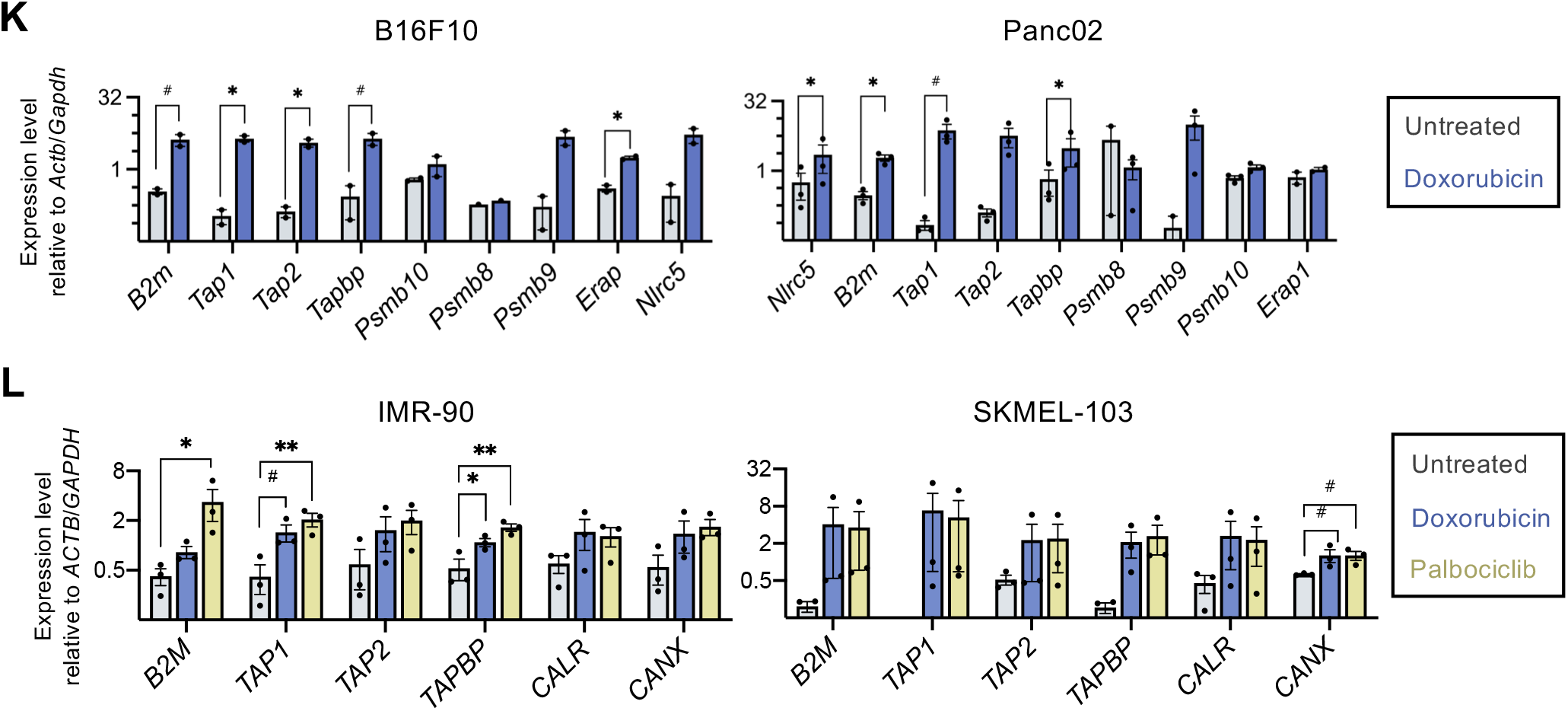
Senescent cells upregulate MHC class I antigen presentation. A. Senescence-associated beta-galactosidase staining of human (SKMEL-103, IMR-90) and murine (B16F10, mouse embryonic fibroblast [MEF]) cells, untreated or exposed to various senescence-inducing stimuli. B. mRNA expression levels of *CDKN1A* in untreated human (SKMEL-103, IMR-90) and *Cdkn2a* murine (B16F10, MEFs) cells, untreated or exposed to various senescence-inducing stimuli of *n*=2-3 independent experiments as indicated. *p<0.05, **p<0.01, Unpaired Student’s t test, compared to untreated cells. C. Flow cytometry analysis of HLA-A/B/C expression in untreated *versus* senescent IMR-90 and SKMEL-103, in which senescence was induced by doxorubicin or palbociclib as indicated, showing the fluorescence signal of each stained sample and its unstained control (lighter color). D. Flow cytometry analysis of H-2Qa1 and H-2Qa2 expression in untreated *versus* senescent MEFs, in which senescence was induced by doxorubicin, nutlin or by late passaging. Representative histograms (left panel) showing the fluorescence signal of each stained sample and its unstained control (lighter color) and quantification after autofluorescence subtraction (right panel) of *n*=3 independent experiments are shown. ***p< 0.001; **p<0.01, one-way ANOVA, compared to untreated MEFs. E. Top upregulated Broad Hallmarks from the differential expression analysis (RNAseq) of senescent Panc02, in which senescence was induced by doxorubicin compared to untreated cells. *n=4* independent biological replicates were analyzed. F. Gene set enrichment analysis (GSEA) of IFN-⍺ and IFN-ɣ response (Broad Hallmarks) genes found upregulated in senescent Panc02 compared to untreated cells. G. Top upregulated Broad Hallmarks from the differential expression analysis (RNAseq) of senescent SK-MEL103, in which senescence was induced by palbociblib as compared to untreated cells. *n=*3 independent biological replicates were analyzed. H. Gene set enrichment analysis (GSEA) of IFN-⍺ and IFN-ɣ response (Broad Hallmarks) genes found upregulated in senescent SKMEL-103 as compared to untreated cells. I. mRNA expression levels of antigen presentation machinery- and immunoproteasome-related genes in untreated *versus* senescent murine cancer B16F10 and Panc02 cell lines measured by qRT-PCR (relative to the average expression of housekeeping genes *Actb* and *Gapdh*). *n*=2-3 independent experiments. *p<0.05, ^#^p<0.1; unpaired Student’s t test, compared to untreated cells. J. RNA expression levels of antigen presentation machinery-related genes in untreated *versus* senescent human SKMEL-103 and IMR-90 cells measured by qRT-PCR (relative to the average expression of housekeeping genes *ACTB* and *GAPDH*). *n*=3 independent experiments. **p<0.01, *p<0.05, ^#^p<0.1; one-way ANOVA test, compared to untreated cells.

**Supplementary Figure S2:**
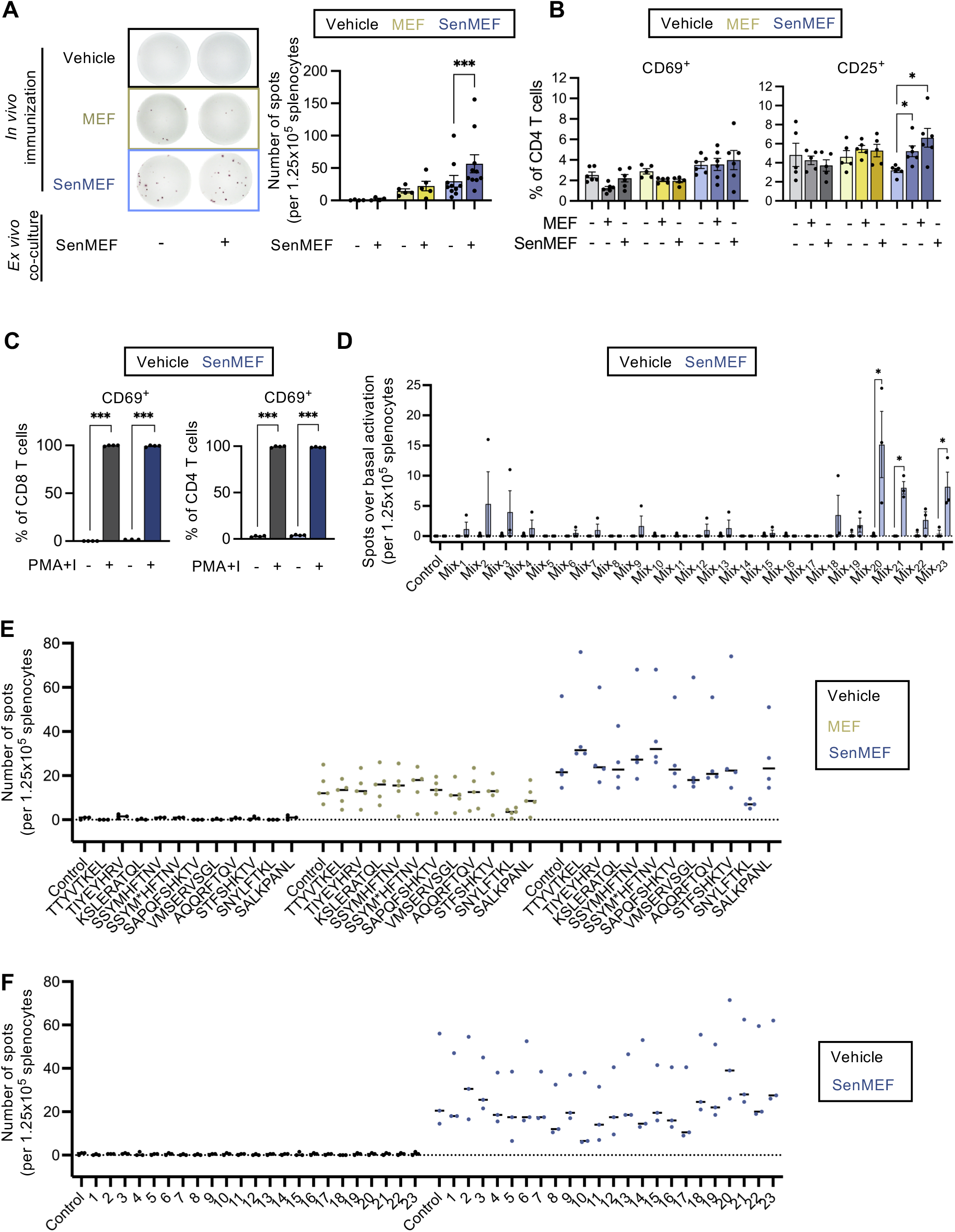
Senescent cells induce an adaptive immune response *in vivo* and present an altered immunopeptidome. A. ELISpot assay to detect IFN-γ production in splenocytes isolated from non-immunized mice or animals immunized with senMEF (*n*=4 mice per group). 2.5×10^5^ or 1.25×10^5^ splenocytes were cultured in RPMI alone or with senFB (1:10 target-to-splenocyte ratio). Representative picture (left panel) and quantification (right panel) are shown. **p<0.01, *p<0.05; unpaired Student’s t test, compared to RPMI alone treatment. B. Measurement of CD4 T cell activation, as monitored via CD69 expression (left panel) or CD25 expression (right panel) in naïve *versus* MEF or senMEF-immunized animals, after coculture with MEF or senMEF *ex vivo* (*n*=4 mice per group). C. Measurement of CD8 (left panel) and CD4 (right panel) T cells activation, as monitored via CD69 expression upon PMA+Ionomycin treatment, used as a positive control of T cell activation. D. Pool of peptides validated using ELISpot assay to detect IFN-γ production in splenocytes isolated from non-immunized mice or animals immunized with senMEF (*n*=3 mice per group). Splenocytes were cultured in RPMI, either alone (control) or supplemented with the remaining peptides obtained from the immunopeptidome analysis pooled in combinations of 1-3 peptides as indicated. The number of spots for each condition above the control condition (background) was quantified and represented. *p<0.05; unpaired Student’s t test, compared to RPMI alone treatment. E. Raw number of spots from ELISpot assay to detect IFN-γ production in splenocytes isolated from non-immunized mice, untreated MEF or animals immunized with senMEF (*n*=3-6 mice per group). Splenocytes were cultured in RPMI, either alone (control), or supplemented with the selected different peptides obtained from the immunopeptidome analysis (as indicated). F. Raw number of spots from ELISpot assay to detect IFN-γ production in splenocytes isolated from non-immunized mice or animals immunized with senMEF (*n*=3 mice per group). Splenocytes were cultured in RPMI, either alone peptide (control) or supplemented with the different peptides pools obtained from the immunopeptidome analysis (as indicated).

**Supplementary Figure S3:**
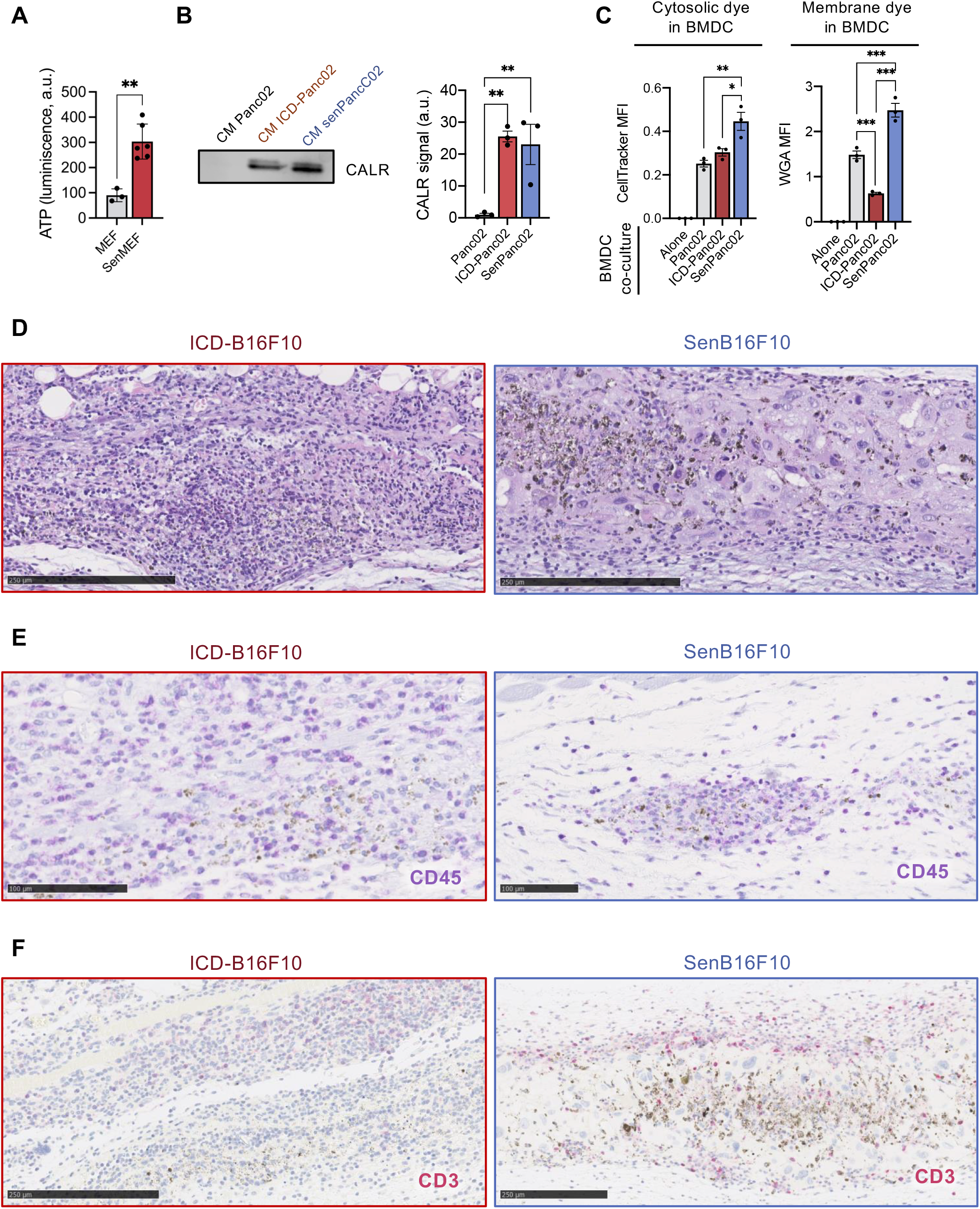
Senescent cancer cells display high immunogenic properties and sustained persistence *in vivo*. A. Levels of extracellular ATP in the conditioned medium (CM) of 10^6^ untreated MEF or senescent MEF induced by a low dose of doxorubicin (SenMEF). *n*=3 independent experiments. **p<0.01 *p<0.05; unpaired Student’s t test. B. Immunoblot detection of CALR in the CM of 10^6^ untreated Panc02 cells, dying by immunogenic cell death induced by a high dose of doxorubicin (ICD-Panc02) or senescent Panc02 induced by a low dose of doxorubicin (senPanc02). Representative image (left panel) and quantification (right panel) of *n*=3 independent experiments are shown. **p<0.01; one-way ANOVA test compared to untreated Panc02 cells. C. Hematoxylin-eosin in skin sections of animals after 7 days of subcutaneous injection of ICD or senB16F10. Note that the brown pigmentation is due to the melanine. Representative images selected by a histopathologist of *n*=5 animals per group are shown. Scale bars for each images are shown (250µm). D. Immunohistochemistry of CD45^+^ cells (purple) in skin sections of animals after 7 days of subcutaneous injection of ICD or senB16F10. Note that the brown pigmentation is due to the melanine. Representative images selected by a histopathologist of *n*=5 animals per group are shown. Scale bars for each images are shown (100µm). F. Immunohistochemistry of CD3^+^ cells (purple) in skin sections of animals 7 days after subcutaneous injection of ICD or senB16F10. Note that the brown pigmentation is due to the melanine. Representative images selected by a histopathologist of *n*=5 animals per group are shown. Scale bars for each images are shown (250µm).

**Supplementary Figure S4:**
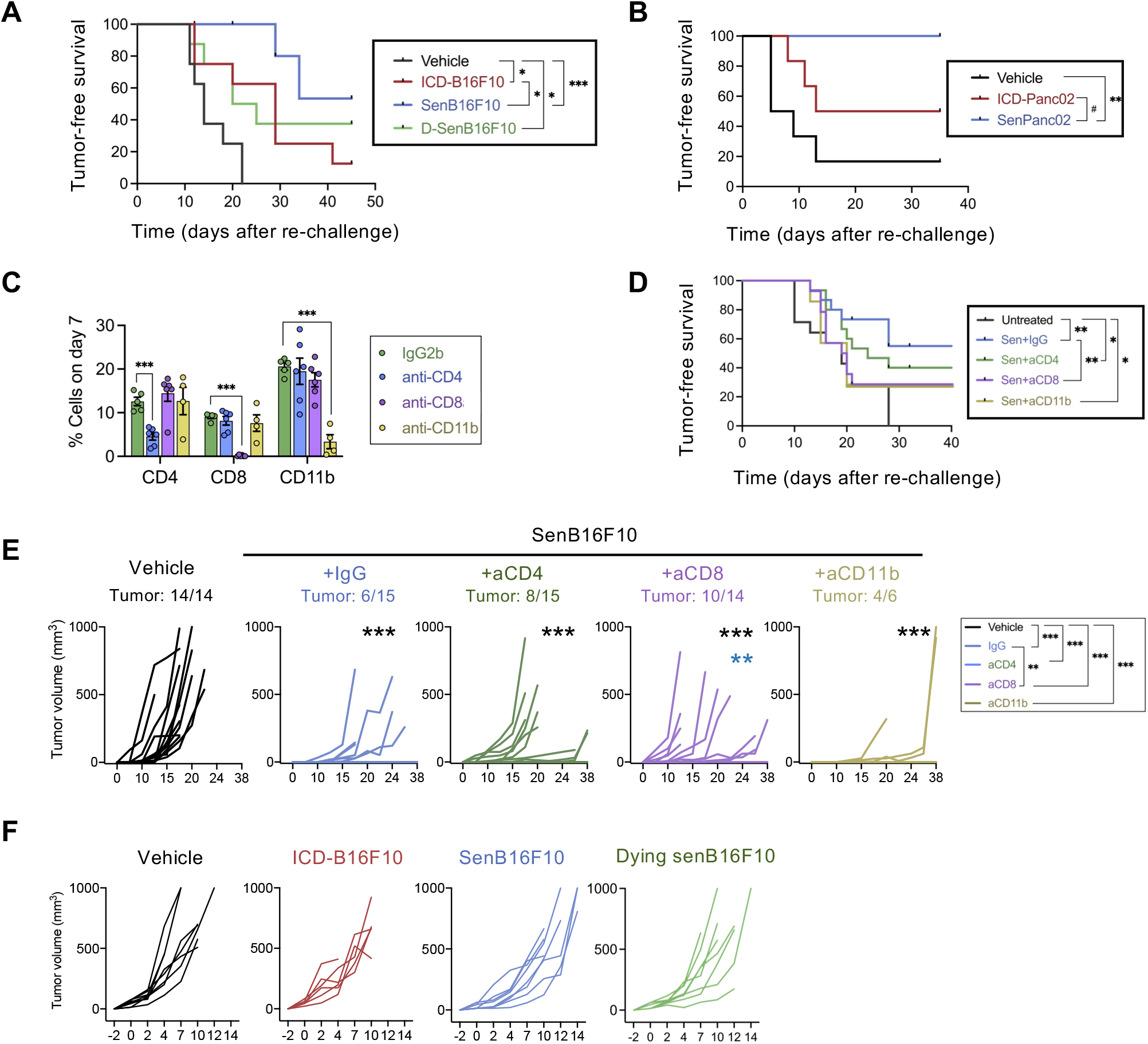
Immunization with senescent cancer cells promotes anti-cancer immune surveillance. A. Tumor-free survival in vehicle-treated mice or mice immunized with B16F10 cells dying by immunogenic cell death induced by a high dose of doxorubicin (ICD-B16F10), senescent B16F10 induced by a low dose of doxorubicin (senB16F10) or senescent B16F10 cells dying by senolysis induced by navitoclax (dying-senB16F10) (*n*=8 mice per group). **p<0.01, *p<0.05; Log-rank (Mantel-Cox) test. B. Tumor-free survival of vehicle-treated mice or mice immunized with PANC02 cells dying by immunogenic cell death induced by a high dose of doxorubicin (ICD-Panc02) or senescent Panc02 induced by a low dose of doxorubicin (senPanc02) (*n*=6 mice per group). **p<0.01, *p<0.05; Log-rank (Mantel-Cox) test. C. Flow cytometry analysis of immune populations (CD4^+^, CD8^+^ and CD11b^+^ cells) in peripheral blood from animals treated with IgG or the indicated blocking antibodies after 7 days of immune depletions (*n*=4-6 mice per group). D. Tumor-free survival of vehicle-treated mice (*n*=14) or mice immunized with senB16F10 treated with IgG (*n*=14) or the indicated blocking antibodies as described in Fig. 4D (*n*=15 for aCD4, *n*=14 for aCD8, or *n*=6 for aCD11b). **p<0.01, *p<0.05; Log-rank (Mantel-Cox) test. E. Individual tumor growth curves from vehicle-treated mice (*n*=14) or mice immunized with senB16F10 treated with IgG (*n*=14) or the indicated blocking antibodies as described in Fig. 4D (*n*=15 for aCD4, *n*=14 for aCD8, or *n*=6 for aCD11b). **p<0.01, *p<0.05; Two-way ANOVA test. F. Individual tumor growth of B16F10 tumor-bearing animals immunized with ICD-B16F10, dying senB16F10 or senB16F10. *p<0.05; Two-way ANOVA test (*n*=7-8).

**Supplementary Figure S5:**
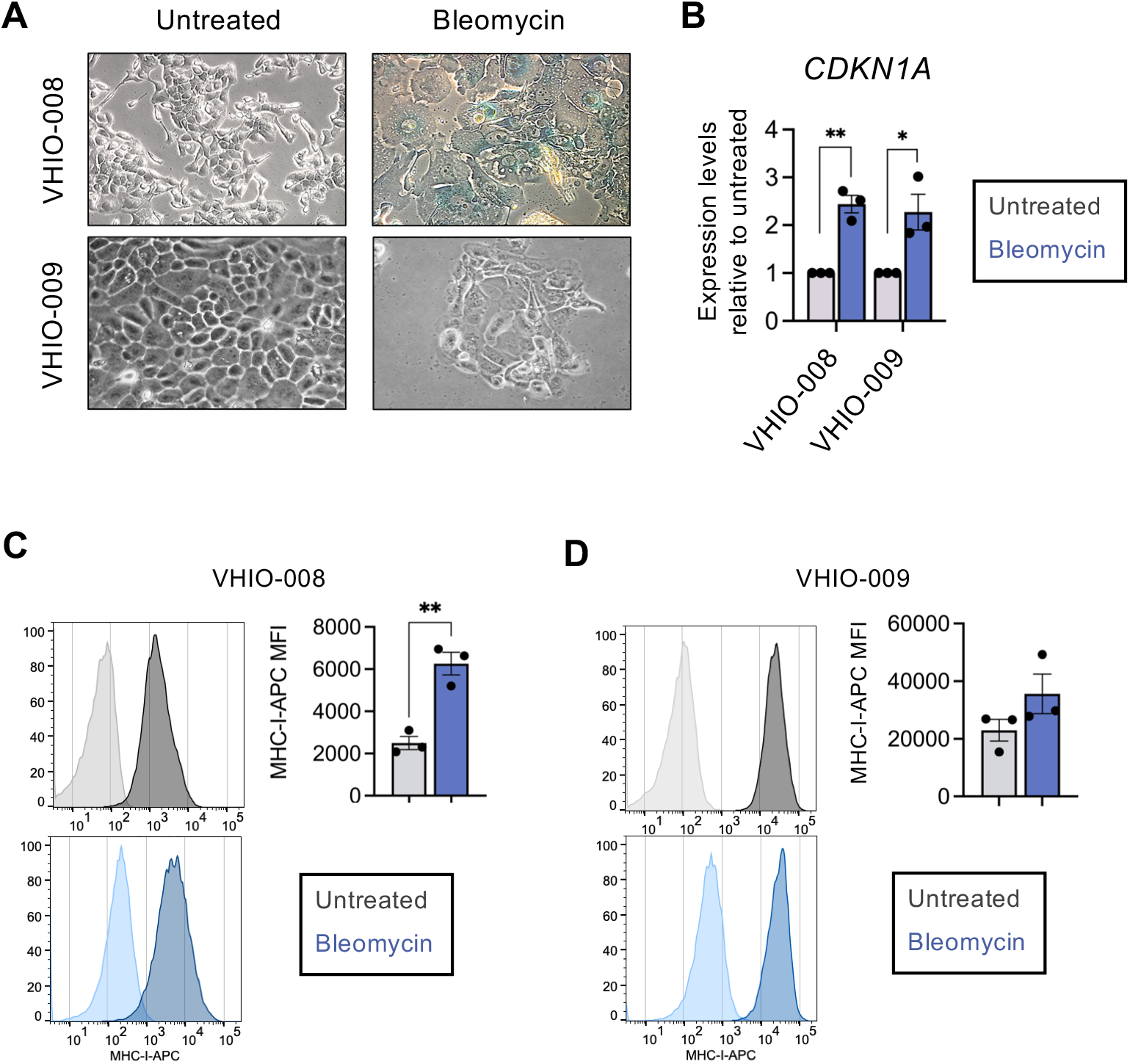
Senescent cancer cells hyper-stimulate autologous reactive TILs from human patients. A. Senescence-associated beta-galactosidase staining of patient-derived VHIO-008 and VHIO-009 cells, untreated or senescent after treatment with bleomycin. B. mRNA expression levels of *CDKN1A* in patient-derived VHIO-008 and VHIO-009 cells, untreated or senescent after treatment with bleomycin of *n=*3 independent experiments as indicated. Relative to untreated cells. *p<0.05, **p<0.01; unpaired Student’s t test, compared to untreated cells. C. Flow cytometry analysis of MHC-I expression in untreated or senescent VHIO-008, treated with bleomycin. Representative histograms (left panel) showing the fluorescence signal of each stained sample and its unstained control (lighter color) and quantification after autofluorescence subtraction (right panel) of *n*=3 independent experiments are shown. **p<0.01, unpaired Student’s t test, compared to untreated cells. D. Flow cytometry analysis of MHC-I expression in untreated or senescent VHIO-009, treated with bleomycin. Representative histograms (left panel) showing the fluorescence signal of each stained sample and its unstained control (lighter color) and quantification after autofluorescence subtraction (right panel) of *n*=3 independent experiments are shown.

## REFERENCES

1. Muñoz-Espín D, Serrano M. Cellular senescence: From physiology to pathology. Nat Rev Mol Cell Biol. Nature Publishing Group; 2014;15:482–96.

2. Coppé JP, Desprez PY, Krtolica A, Campisi J. The senescence-associated secretory phenotype: The dark side of tumor suppression. Annu Rev Pathol Mech Dis. 2010;5:99–118.

3. Birch J, Gil J. Senescence and the SASP: Many therapeutic avenues. Genes Dev. 2020;34:1565–76.

4. Prata LGPL, Ovsyannikova IG, Tchkonia T, Kirkland JL. Senescent cell clearance by the immune system: Emerging therapeutic opportunities. Semin Immunol. Elsevier; 2018;40:101275.

5. Song P, An J, Zou M-H. Immune Clearance of Senescent Cells to Combat Ageing and Chronic Diseases. Cells. 2020;9:671.

6. Kale A, Sharma A, Stolzing A, Stolzing A, Desprez PY, Desprez PY, et al. Role of immune cells in the removal of deleterious senescent cells. Immun Ageing. Immunity & Ageing; 2020;17:1–9.

7. Kang TW, Yevsa T, Woller N, Hoenicke L, Wuestefeld T, Dauch D, et al. Senescence surveillance of pre-malignant hepatocytes limits liver cancer development. Nature. Nature Publishing Group; 2011;479:547–51.

8. Muñoz-Espín D, Cañamero M, Maraver A, Gómez-López G, Contreras J, Murillo-Cuesta S, et al. Programmed cell senescence during mammalian embryonic development. Cell. 2013;155:1104.

9. Egashira M, Hirota Y, Shimizu-Hirota R, Saito-Fujita T, Haraguchi H, Matsumoto L, et al. F4/80+ macrophages contribute to clearance of senescent cells in the mouse postpartum uterus. Endocrinology. 2017;158:2344–53.

10. Xue W, Zender L, Miething C, Dickins RA, Hernando E, Krizhanovsky V, et al. Senescence and tumour clearance is triggered by p53 restoration in murine liver carcinomas. Nature. 2007;445:656–60.

11. Sagiv A, Burton DGA, Moshayev Z, Vadai E, Wensveen F, Ben-Dor S, et al. NKG2D ligands mediate immunosurveillance of senescent cells. Aging (Albany NY). 2016;8:328–44.

12. Brighton PJ, Maruyama Y, Fishwick K, Vrljicak P, Tewary S, Fujihara R, et al. Clearance of senescent decidual cells by uterine natural killer cells in cycling human endometrium. Elife. 2017;6:1–23.

13. Ovadya Y, Landsberger T, Leins H, Vadai E, Gal H, Biran A, et al. Impaired immune surveillance accelerates accumulation of senescent cells and aging. Nat Commun. Springer US; 2018;9.

14. Pereira BI, Devine OP, Vukmanovic-Stejic M, Chambers ES, Subramanian P, Patel N, et al. Senescent cells evade immune clearance via HLA-E-mediated NK and CD8+ T cell inhibition. Nat Commun. Springer US; 2019;10.

15. Krizhanovsky V, Yon M, Dickins RA, Hearn S, Simon J, Miething C, et al. Senescence of Activated Stellate Cells Limits Liver Fibrosis. Cell. 2008;134:657–67.

16. Iannello A, Thompson TW, Ardolino M, Lowe SW, Raulet DH. p53-dependent chemokine production by senescent tumor cells supports NKG2D-dependent tumor elimination by natural killer cells. J Exp Med. 2013;210:2057–69.

17. Arora S, Thompson PJ, Wang Y, Bhattacharyya A, Apostolopoulou H, Hatano R, et al. Invariant natural killer T cells coordinate removal of senescent cells. Med. Elsevier Inc.; 2021;2:938–950.e8.

18. Binet F, Cagnone G, Crespo-Garcia S, Hata M, Neault M, Dejda A, et al. Neutrophil extracellular traps target senescent vasculature for tissue remodeling in retinopathy. Science (80). 2020;369.

19. Frescas D, Roux CM, Aygun-Sunar S, Gleiberman AS, Krasnov P, Kurnasov O V., et al. Senescent cells expose and secrete an oxidized form of membrane-bound vimentin as revealed by a natural polyreactive antibody. Proc Natl Acad Sci U S A. 2017;114:E1668–77.

20. van Tuyn J, Jaber-Hijazi F, MacKenzie D, Cole JJ, Mann E, Pawlikowski JS, et al. Oncogene-Expressing Senescent Melanocytes Up-Regulate MHC Class II, a Candidate Melanoma Suppressor Function. J Invest Dermatol. 2017;137:2197–207.

21. Raval A, Puri N, Rath PC, Saxena RK. Cytokine regulation of expression of class I MHC antigens. Exp Mol Med. 1998;30:1–13.

22. Frisch SM, MacFawn IP. Type I interferons and related pathways in cell senescence. Aging Cell. 2020;19:1–12.

23. Kobayashi KS, Van Den Elsen PJ. NLRC5: A key regulator of MHC class I-dependent immune responses. Nat Rev Immunol. Nature Publishing Group; 2012;12:813–20.

24. Reits EA, Hodge JW, Herberts CA, Groothuis TA, Chakraborty M, Wansley EK, et al. Radiation modulates the peptide repertoire, enhances MHC class I expression, and induces successful antitumor immunotherapy. J Exp Med. 2006;203:1259–71.

25. Gravett AM, Trautwein N, Stevanović S, Dalgleish AG, Copier J. Gemcitabine alters the proteasome composition and immunopeptidome of tumour cells. Oncoimmunology. 2018;7.

26. Oh CY, Klatt MG, Bourne C, Dao T, Dacek MM, Brea EJ, et al. ALK and RET inhibitors promote HLA Class i antigen presentation and unmask new antigens within the tumor immunopeptidome. Cancer Immunol Res. 2019;7:1984–97.

27. Stopfer LE, Mesfin JM, Joughin BA, Lauffenburger DA, White FM. Multiplexed relative and absolute quantitative immunopeptidomics reveals MHC I repertoire alterations induced by CDK4/6 inhibition. Nat Commun. 2020;11:1–14.

28. Starck SR, Tsai JC, Chen K, Shodiya M, Wang L, Yahiro K, et al. Translation from the 5’ untranslated region shapes the integrated stress response. Science (80). 2016;351.

29. Schuster H, Shao W, Weiss T, Pedrioli PGA, Roth P, Weller M, et al. A tissue-based draft map of the murine MHC class I immunopeptidome. Sci Data. 2018;5:1–11.

30. Mellman I, Steinman RM. Dendritic cells: Specialized and regulated antigen processing machines. Cell. 2001;106:255–8.

31. Jhunjhunwala S, Hammer C, Delamarre L. Antigen presentation in cancer: insights into tumour immunogenicity and immune evasion. Nat Rev Cancer. Springer US; 2021;21:298–312.

32. Basisty N, Kale A, Jeon O, Kuehnemann C, Payne T, Rao C, et al. A Proteomic Atlas of Senescence-Associated Secretomes for Aging Biomarker Development. SSRN Electron J. 2019;1–26.

33. Galluzzi L, Vitale I, Warren S, Adjemian S, Agostinis P, Martinez AB, et al. Consensus guidelines for the definition, detection and interpretation of immunogenic cell death. J Immunother Cancer. 2020;8:1–22.

34. Kroemer G, Galassi C, Zitvogel L, Galluzzi L. Immunogenic cell stress and death. Nat Immunol. Springer US; 2022;23:487–500.

35. Mohapatra A Das, Tirrell I, Benechet AP, Pattnayak S, Khanna KM, Srivastava PK. Cross-dressing of CD8aþ dendritic cells with antigens from live mouse tumor cells is a major mechanism of cross-priming. Cancer Immunol Res. 2020;8:1287–99.

36. Wang J, Saffold S, Cao X, Krauss J, Chen W. Eliciting T cell immunity against poorly immunogenic tumors by immunization with dendritic cell-tumor fusion vaccines. J Immunol. 1998;161:5516–24.

37. Zitvogel L, Perreault C, Finn OJ, Kroemer G. Beneficial autoimmunity improves cancer prognosis. Nat Rev Clin Oncol. Springer US; 2021;18:591–602.

38. Caron E, Vincent K, Fortier MH, Laverdure JP, Bramoullé A, Hardy MP, et al. The MHC i immunopeptidome conveys to the cell surface an integrative view of cellular regulation. Mol Syst Biol. 2011;7:1–15.

39. Serrano M, Lin AW, McCurrach ME, Beach D, Lowe SW. Oncogenic ras provokes premature cell senescence associated with accumulation of p53 and p16(INK4a). Cell. 1997;88:593–602.

40. Bø TH, Dysvik B, Jonassen I. LSimpute: accurate estimation of missing values in microarray data with least squares methods. Nucleic Acids Res. 2004;32:e34.

41. Eden E, Navon R, Steinfeld I, Lipson D, Yakhini Z. GOrilla: A tool for discovery and visualization of enriched GO terms in ranked gene lists. BMC Bioinformatics. 2009;10:1–7.

42. Martin M. Cutadapt removes adapter sequences from high-throughput sequencing reads. , may. MBnet.journal. 2011;17:10–2.

43. Dobin A, Davis CA, Schlesinger F, Drenkow J, Zaleski C, Jha S, et al. STAR: ultrafast universal RNA-seq aligner. Bioinformatics. 2013;29:15–21.

44. Tarasov A, Vilella AJ, Cuppen E, Nijman IJ, Prins P. Sambamba: fast processing of NGS alignment formats. Bioinformatics. 2015;31:2032–4.

45. R Core Team. R: A language and environment for statistical computing. 2017;

46. Rossell D, Stephan-Otto Attolini C, Kroiss M, Stöcker A. Quantifying alternative splicing from paired-end rna-sequencing data. Ann Appl Stat. 2014;8:309–30.

47. Ritchie ME, Phipson B, Wu D, Hu Y, Law CW, Shi W, et al. limma powers differential expression analyses for RNA-sequencing and microarray studies. Nucleic Acids Res. 2015;43:e47.

48. Liberzon A, Birger C, Thorvaldsdóttir H, Ghandi M, Mesirov JP, Tamayo P. The Molecular Signatures Database (MSigDB) hallmark gene set collection. Cell Syst. 2015;1:417–25.

49. Carlson M. org.Hs.eg.db: Genome wide annotation for Human. R package version 3.7.0.

50. Wu D, Lim E, Vaillant F, Asselin-Labat M-L, Visvader JE, Smyth GK. ROAST: rotation gene set tests for complex microarray experiments. Bioinformatics. 2010;26:2176–82.

51. Efron B RT. On testing the significance of sets of genes. Ann Appl Stat. 2007;1:107–29.

52. Caron E, Kowalewski DJ, Chiek Koh C, Sturm T, Schuster H, Aebersold R. Analysis of Major Histocompatibility Complex (MHC) Immunopeptidomes Using Mass Spectrometry. Mol Cell Proteomics. 2015;14:3105–17.

53. Sirois I, Isabelle M, Duquette JD, Saab F, Caron E. Immunopeptidomics: Isolation of Mouse and Human MHC Class I- and II-Associated Peptides for Mass Spectrometry Analysis. J Vis Exp. United States; 2021;

54. Kovalchik KA, Ma Q, Wessling L, Saab F, Duquette JD, Kubiniok P, et al. MhcVizPipe: A Quality Control Software for Rapid Assessment of Small- to Large-Scale Immunopeptidome Datasets. Mol Cell Proteomics. 2022;21:100178.

55. Oliveros JC. VENNY. An interactive tool for comparing lists with Venn Diagrams. 2007; Available from: https://bioinfogp.cnb.csic.es/tools/venny/index.html

56. Bankhead P, Loughrey MB, Fernández JA, Dombrowski Y, McArt DG, Dunne PD, et al. QuPath: Open source software for digital pathology image analysis. Sci Rep. 2017;7:1–7.

57. Tran E, Ahmadzadeh M, Lu Y-C, Gros A, Turcotte S, Robbins PF, et al. Immunogenicity of somatic mutations in human gastrointestinal cancers. Science. 2015;350:1387–90.

